# Hap2-Ino80 facilitated transcription promotes *de novo* establishment of CENP-A chromatin

**DOI:** 10.1101/778639

**Authors:** Puneet P. Singh, Manu Shukla, Sharon A. White, Pin Tong, Tatsiana Auchynnikava, Christos Spanos, Juri Rappsilber, Alison L. Pidoux, Robin C. Allshire

## Abstract

Centromeres are maintained epigenetically by the presence of CENP-A, an evolutionarily-conserved histone H3 variant, which directs kinetochore assembly and hence, centromere function. To identify factors that promote assembly of CENP-A chromatin, we affinity selected solubilised fission yeast CENP-A^Cnp1^ chromatin. All subunits of the Ino80 complex were enriched, including the auxiliary subunit Hap2. In addition to a role in maintenance of CENP-A^Cnp1^ chromatin integrity at endogenous centromeres, Hap2 is required for *de novo* assembly of CENP-A^Cnp1^ chromatin on naïve centromere DNA and promotes H3 turnover on centromere regions and other loci prone to CENP-A^Cnp1^ deposition. Prior to CENP-A^Cnp1^ chromatin assembly, Hap2 facilitates transcription from centromere DNA. These analyses suggest that Hap2-Ino80 destabilises H3 nucleosomes on centromere DNA through transcription-coupled histone H3 turnover, driving the replacement of resident H3 nucleosomes with CENP-A^Cnp1^ nucleosomes. These inherent properties define centromere DNA by directing a program that mediates CENP-A^Cnp1^ assembly on appropriate sequences.

## INTRODUCTION

The accurate delivery of all chromosomes to both resulting nuclei during mitotic cell division is required for eukaryotic cell viability and to prevent aneuploidy, a hallmark of cancer (Kops et al., 2005). The centromere region of chromosomes mediates their attachment to spindle microtubules for normal mitotic chromosome segregation (Cleveland et al., 2003; Fukagawa and Earnshaw, 2014; Pidoux and Allshire, 2004). In many organisms, centromeres are assembled on repetitive elements such as α-satellite repeats, minor satellite repeats, cen180/CentC/CentO repeats, and retroelements at human, mouse, plant, and *Drosophila* centromeres, respectively (Chang et al., 2019; Cheng et al., 2002; Grady et al., 1992; Kipling et al., 1991). Although such centromere repeats lack sequence similarity, in many cases their introduction as naked DNA into cells triggers *de novo* kinetochore assembly (Allshire and Karpen, 2008; McKinley and Cheeseman, 2016; Müller and Almouzni, 2017).

The underlying conserved feature at eukaryotic centromeres is the assembly of nucleosomes containing the histone H3 variant CENP-A (also generally known as cenH3; and specifically as CID in *Drosophila*, Cse4 in *Saccharomyces cerevisiae*, Cnp1 in *Schizosaccharomyces pombe*) in place of canonical H3 to direct kinetochore assembly on such repetitive elements. Moreover, it is known that following deletion of an endogenous centromere, CENP-A incorporation and neocentromeres can arise at novel non-centromeric DNA locations (Ishii et al., 2008; Ketel et al., 2009; Shang et al., 2013). CENP-A chromatin has also been shown to be sufficient to trigger kinetochore assembly (Barnhart et al., 2011; Chen et al., 2014; Hori et al., 2013; Mendiburo et al., 2011). Thus, CENP-A deposition and not the primary sequence of centromere DNA determines the position of centromere formation. However, centromere DNA itself may harbour properties that favour CENP-A and kinetochore assembly (Baum et al., 1994; Fachinetti et al., 2015; Logsdon et al., 2019).

Three fission yeast species possess complex regional centromeres in which CENP-A^Cnp1^ and kinetochores are assembled over a non-repetitive central domain of approximately 10 kb in *S. pombe*, *S. octosporus* and *S. cryophilus* (Tong et al., 2019). Although flanking gene order is preserved, the central domain sequence is not conserved amongst these species. Despite this lack of similarity, central domain DNA from S*. octosporus* and *S. cryophilus,* as well as *S. pombe*, can direct *de novo* CENP-A^Cnp1^ and kinetochore assembly in *S. pombe* (Catania et al., 2015; Folco et al., 2008; Tong et al., 2019). This suggests that these non-homologous centromere DNAs possess innate features that program events which preferentially trigger the assembly of CENP-A^Cnp1^ in place of H3 nucleosomes.

During replication, parental CENP-A has been shown to distribute equally to nucleosomes on duplicated DNA of both sister centromeres, thus halving the amount of CENP-A at each centromere (Black and Cleveland, 2011; Jansen et al., 2007; Schuh et al., 2007). Replenishment by deposition of newly-synthesised CENP-A is temporally separated from DNA replication-coupled H3 chromatin assembly. The timing of new CENP-A deposition differs in various species; mitosis/late telophase/early G1 in human and *Drosophila* (Erhardt et al., 2008; Mellone et al., 2011; Schuh et al., 2007), S phase in *S. cerevisiae* (Jansen et al., 2007; Pearson et al., 2004; Wisniewski et al., 2014), and G2 in plants and *S. pombe* (Lando et al., 2012; Lermontova et al., 2006; Shukla et al., 2018).

Several studies have reported the association of RNA polymerase II (RNAPII) with, and/or transcription from CENP-A-associated DNA so that non-coding transcription has become an apparent integral feature of centromeres (Duda et al., 2017). RNAPII has been detected at human artificial chromosome (HAC; Bergmann et al., 2011), human metaphase (Chan et al., 2012), *Drosophila* (Bobkov et al., 2018), and *S. cerevisiae* (Ohkuni and Kitagawa, 2011) centromeres, and also the central domains of fission yeast centromeres (Catania et al., 2015). Disruption of centromere transcription appears to hinder CENP-A loading and/or maintenance (Bergmann et al., 2011; Cardinale et al., 2009; Chen et al., 2015; Ling and Yuen, 2019; McNulty et al., 2017; Nakano et al., 2008). Moreover, increased centromere transcription results in the rapid loss of CENP-A and centromere function (Bergmann et al., 2012; Hill and Bloom, 1987). The central CENP-A^Cnp1^ domains from *S. pombe* centromeres contain numerous RNAPII transcriptional start sites and promoters (Catania et al., 2015; Choi et al., 2011). In addition, ectopically-located central domain DNA, that lacks CENP-A^Cnp1^ chromatin, exhibits high rates of histone H3 turnover (Shukla et al., 2018). Such observations suggest that transcription-coupled chromatin remodelling events might drive the eviction of H3 and its replacement with CENP-A^Cnp1^. Consistent with this view, the accumulation of elongating RNAPII on ectopic centromere DNA during G2 coincides with the eviction of H3 and deposition of new CENP-A (Shukla et al., 2018). However, it remains to be determined which transcription-associated chromatin remodelling factors provoke the replacement of H3 with CENP-A on naïve centromere DNA.

Various ATP-dependent chromatin remodelling complexes provide access to the underlying DNA of chromatin-coated templates. Their activities enable transcription by disassembling nucleosomes, sliding nucleosomes, or replacing nucleosomal histone subunits with transcription-promoting variants (Becker and Hörz, 2002; Clapier and Cairns, 2009). The Remodelling and Spacing Factor (RSF) interacts with CENP-A chromatin in mid-G1, its depletion reduces CENP-A levels at centromeres (Perpelescu et al., 2009) and tethering of RSF1 to centromere repeats promotes histone turnover/exchange resulting in both H3.3 and CENP-A deposition (Ohzeki et al., 2016). In *S. pombe*, Hrp1 (ortholog of Chromo-Helicase DNA-binding protein 1, CHD1) is enriched at centromeres and is required to maintain normal CENP-A^Cnp1^ levels at centromeres (Choi et al., 2011; Walfridsson et al., 2005). Loss of the histone chaperone FAcilitates Chromatin Transcription (FACT) results in promiscuous CENP-A^Cnp1^ assembly at non-centromeric locations A^Cnp1^ (Choi et al., 2012), suggesting that CENP-A^Cnp1^ may be titrated away from centromeres by loss of such factors. Moreover, inducible ectopic centromeres in *Drosophila* requires FACT-mediated RNAPII-dependent transcription of underlying DNA, indicating a necessity for transcription during CENP-A assembly (Chen et al., 2015).

Ino80 is a Snf2-family ATPase evolutionarily conserved from yeast to humans, that participates in transcription, DNA replication and DNA repair (Bao and Shen, 2007; Conaway and Conaway, 2009; Shen et al., 2000). The Ino80 complex (Ino80C) can slide nucleosomes in an ATP-dependent manner (Chen et al., 2011; Shen et al., 2000; Willhoft et al., 2016) and can space multiple nucleosomes on longer DNA fragments (Udugama et al., 2011). Ino80C may also remove H2A.Z–H2B dimers from nucleosomes, replacing them with H2A–H2B dimers (Brahma et al., 2017; Papamichos-Chronakis et al., 2011). H2A.Z is enriched in +1 nucleosomes downstream of promoters of many active genes and loss of Ino80 function affects transcription (Gai et al., 2017; Hogan et al., 2010; Klopf et al., 2009; Xue et al., 2015). Individual Ino80C subunits make up three modules that associate with the main Ino80 ATPase subunit (Table S1; Chen et al., 2011; Tosi et al., 2013; Watanabe et al., 2015). *S. pombe* Ino80 has been shown to influence the maintenance of CENP-A^Cnp1^ chromatin at centromeres (Choi et al., 2017). However, it is not known if Ino80C influences centromere DNA transcription or the establishment of CENP-A chromatin on naïve centromere DNA and thus centromere identity.

Here we utilize affinity selection of CENP-A^Cnp1^ chromatin and mass spectrometry to identify proteins enriched in CENP-A^Cnp1^ chromatin that may promote CENP-A^Cnp1^ assembly. We identify Hap2 as an auxiliary subunit of Ino80C that is required for the conversion of H3 chromatin to CENP-A^Cnp1^ chromatin on naïve centromere DNA. We show that loss of Hap2 function reduces transcription and histone H3 turnover on centromere DNA sequences. Our findings indicate that Hap2-Ino80 is required to promote transcription-associated chromatin remodelling events that drive H3 nucleosome eviction and the assembly of CENP-A^Cnp1^ nucleosomes in their place.

## RESULTS

### Hap2 is an Ino80C subunit that is enriched in CENP-A^Cnp1^ chromatin

To identify proteins involved in the assembly of CENP-A^Cnp1^ chromatin, GFP-tagged CENP-A^Cnp1^ chromatin was affinity purified from micrococcal nuclease (MNase) solubilized chromatin extracts (Figure 1A). Quantitative PCR (qPCR) analysis of the resulting enriched native chromatin revealed significant enrichment of centromeric DNA from the single copy central CENP-A^Cnp1^ domain of cen2 (cc2) compared to the 18 copies of flanking outer repeat (*dg*) sequences (Figure S1A). In addition, SDS-PAGE analysis showed core histones to be prevalent in this affinity-selected GFP-CENP-A^Cnp1^ material (Figure 1B and S1B). Label-free quantitative mass spectrometry detected strong enrichment of all known subunits of both the inner and outer kinetochore complexes (Figure 1C; Table S2). As our procedure also showed association of the chaperones Scm3 (HJURP) and Sim3 (NASP) that mediate CENP-A^Cnp1^ deposition we reasoned that other enriched, but non-centromere specific, proteins might be involved in the incorporation of CENP-A^Cnp1^ into centromeric chromatin.

**Figure 1.**
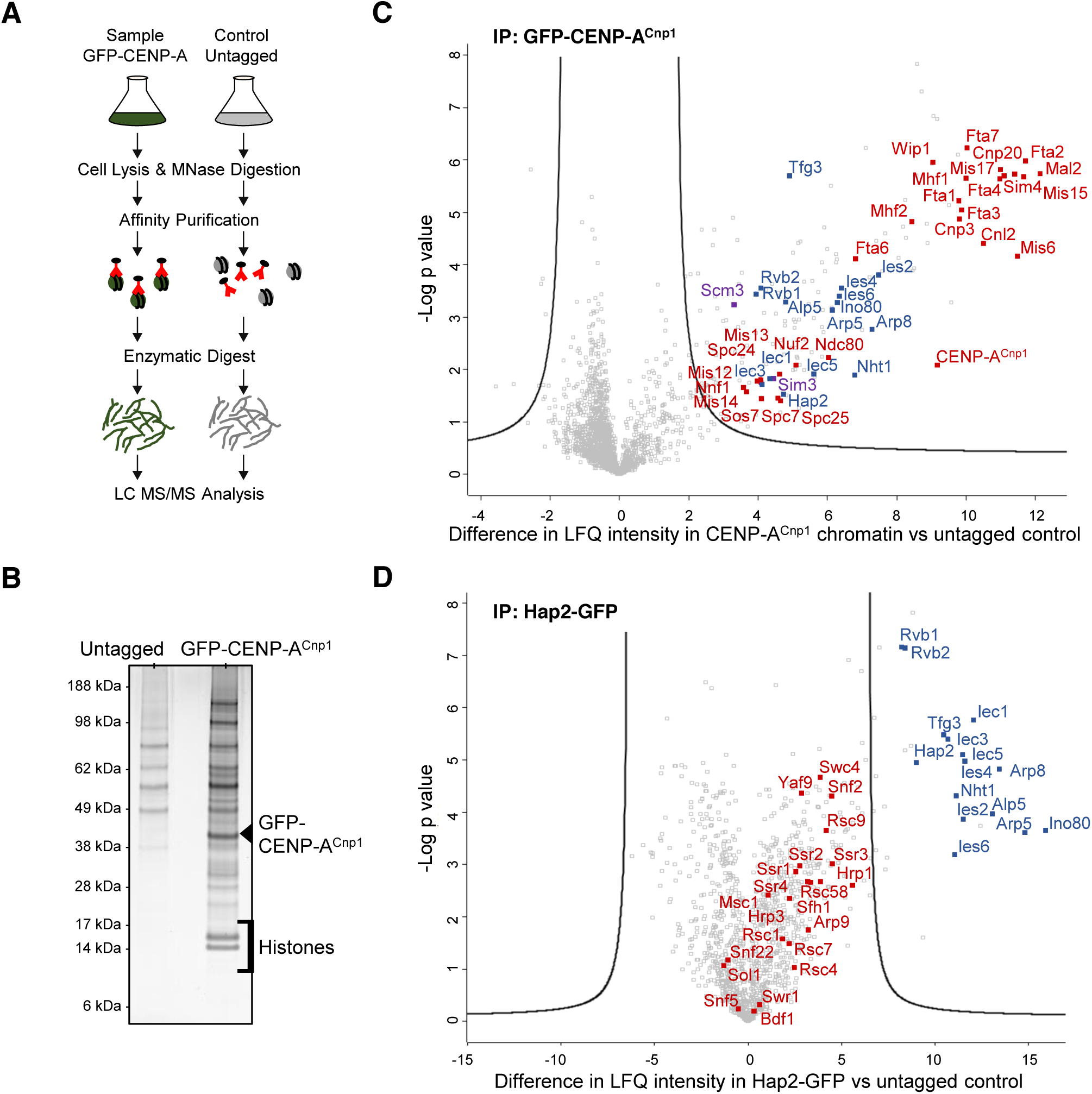
Hap2 is associated with CENP-A^Cnp1^ chromatin and is a subunit of Ino80 complex. (A) Scheme for enrichment of CENP-A^Cnp1^ chromatin associated proteins and identification by mass spectrometry. (B) Silver-stained SDS-PAGE of proteins enriched in IPs from GFP-CENP-A^Cnp1^ or untagged control cells. (C) Volcano plot comparing label free quantification (LFQ) intensity of proteins enriched in affinity selected GFP-CENP-A^Cnp1^ (anti-GFP) versus untagged control. Inner and outer kinetochore proteins detected (Red). Ino80 complex subunits (Blue). CENP-A^Cnp1^ chaperones Scm3 and Sim3 (Purple). (D) Volcano plot comparing label free quantification of proteins enriched in affinity selected Hap2-GFP (anti-GFP) versus untagged control. Ino80 complex subunits (Blue). Swr1 complex subunits, SWI/SNF complex, RSC complex and CHD family members (Red).

It was notable that all subunits of the Ino80 chromatin remodelling complex (Ino80C) were enriched in our affinity-selected GFP-CENP-A^Cnp1^ preparations (Figure 1C). Also enriched in these samples was the low molecular weight alpha-helical Hap2 protein. Hap2 has previously been reported to associate with Ino80C but no characterisation has been performed (Hogan et al., 2010). To confirm association of Hap2 with Ino80C, Hap2 was C-terminally tagged with GFP at its endogenous chromosomal locus (Hap2-GFP; Figure S2A). Affinity selection of Hap2-GFP followed by proteomic analysis revealed that all subunits of Ino80C, but not subunits of other remodelling complexes, are enriched with Hap2-GFP (Figure 1D). Western analysis confirmed that HA-tagged Ino80 (Ino80-HA) associates with immunoprecipitated Hap2-GFP (Figure S1C). Furthermore, affinity selection of Ino80-HA enriched all known Ino80C subunits along with Hap2 (Figure S1D). We conclude that Hap2 is a non-canonical subunit of the fission yeast Ino80 complex and that all Ino80C subunits are enriched in CENP-A^Cnp1^ chromatin.

### Hap2 associates with central domain DNA assembled in CENP-A^Cnp1^ chromatin or H3 chromatin

Microscopic analysis showed that Hap2-GFP localizes in several prominent nuclear foci (Figure 2A). Quantitative chromatin immunoprecipitation (qChIP) assays revealed that Hap2-GFP associates with most chromatin regions analysed including the central CENP-A^Cnp1^ domain of centromeres (cc2), the flanking pericentromeric outer repeats (*dg*) and on the highly expressed *act1^+^* gene (Figure 2B). When 8.5 kb of cen2 central domain DNA (*cen2-cc2*) is inserted at the *ura4* locus on a chromosome arm (*ura4:cc2*), it is assembled in H3 instead of CENP-A^Cnp1^ chromatin (Choi et al., 2012; Shukla et al., 2018). Replacement of 6.5 kb of endogenous *cen2-cc2* DNA with 5.5 kb of *cen1* central domain DNA (*cc2Δ::cc1*) allows analysis across the resulting unique ectopic copy of cen2 central domain DNA (*ura4:cc2*) in the absence CENP-A^Cnp1^ and kinetochore proteins (Figure 2C Top; Choi et al. 2012; Catania et al. 2015; Shukla et al. 2018). qChIP analysis revealed that Hap2-GFP associates with this ectopic *cc2* central domain chromatin (Figure 2C Bottom). A noticeably higher level of Hap2-GFP was consistently detected across ectopic *ura4:cc2* central domain DNA assembled in H3 chromatin relative to the same DNA sequence at the native *cen2* central domain assembled in CENP-A^Cnp1^ chromatin (Figure 2D). Loss of Hap2 does not affect the association of Ino80-HA with these regions (Figure S2B). We conclude that the Ino80C subunit Hap2 is a nuclear protein which associates with non-centromeric loci and is preferentially recruited to centromeric central domain chromatin when assembled in H3 rather than CENP-A^Cnp1^ chromatin.

**Figure 2.**
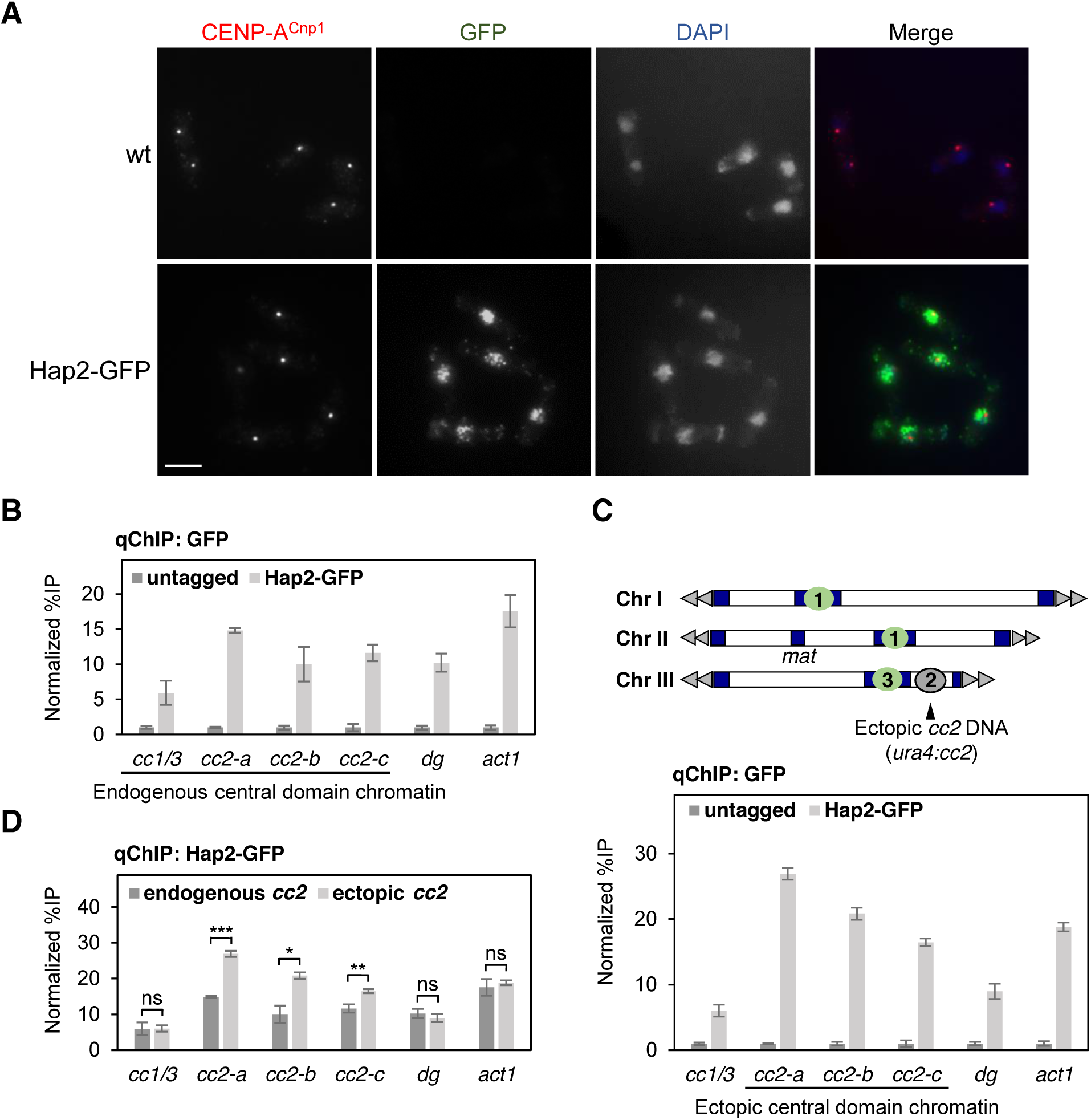
Hap2 is distributed throughout the nucleus and associates with endogenous and ectopic central core chromatin. (A) Immunolocalization of Hap2-GFP in cells. Representative images of wild-type and Hap2-GFP cells stained with anti-CENP-A^Cnp1^ (red), anti-GFP (green) and DAPI (blue). Scale bar, 5 μm. (B) qChIP for Hap2-GFP at four locations within endogenous centromeres (*cc1/3*, *cc2-a*, *cc2-b* and *cc2-c*), outer repeat heterochromatin (*dg*) and non-centromere locus (*act1^+^*). Error bars indicate mean ± SD (n = 3). (C) Top: Diagram indicating strain with endogenous *cen2-cc2* replaced with *cen1* central domain DNA (*cc1*) with insertion of 8.5 kb of *cc2* DNA at the non-centromeric ura4 locus (*ura4:cc2*) on Chr III. Bottom: qChIP for Hap2-GFP at three locations within ectopic central domain H3 chromatin (*cc2-a*, *cc2-b* and *cc2-c*), endogenous centromeres (*cc1/3*), outer repeat heterochromatin (*dg*) and non-centromere locus (*act1^+^*). Error bars indicate mean ± SD (n = 3). (D) Comparison of Hap2-GFP association with central domain sequence assembled in CENP-A^Cnp1^ chromatin or H3 chromatin (data from B and C). Error bars indicate mean ± SD (n = 3). Significance of the differences observed between cells containing endogenous centromeres and ectopic central domain DNA was evaluated using Student’s t-test; * *p* < 0.05, * *p* < 0.005 ***, *p* < 0.005 and n.s., not significant.

### Hap2 is required to maintain CENP-A^Cnp1^ chromatin across endogenous centromeres

Cells lacking Hap2 exhibit an elevated frequency of lagging chromosomes during mitosis, indicating that loss of Hap2 may affect centromere function (Figure 3A). Defective centromere function can result from reduced pericentromeric heterochromatin formation on *dg*/*dh* outer repeats or CENP-A^Cnp1^ chromatin/kinetochore assembly and these can be sensitively detected by the use of silent *ura4^+^* reporter genes inserted within outer repeat heterochromatin or central CENP-A^Cnp1^ domain chromatin (Allshire et al., 1994, 1995; Partridge et al., 2000). No alleviation of heterochromatin-mediated silencing at *otr1:ura4^+^* was detected in *hap2*Δ cells relative to wild-type as indicated by similar poor growth on selective plates lacking uracil (-URA) and good growth on counter-selective 5-FOA plates (Figure 3B and S3A). Consistent with this observation, no significant change was detected in the levels of the heterochromatin H3K9me2 mark or *dg* transcripts produced by the underlying outer repeats in *hap2*Δ relative wild-type cells (Figure 3C and D).

**Figure 3.**
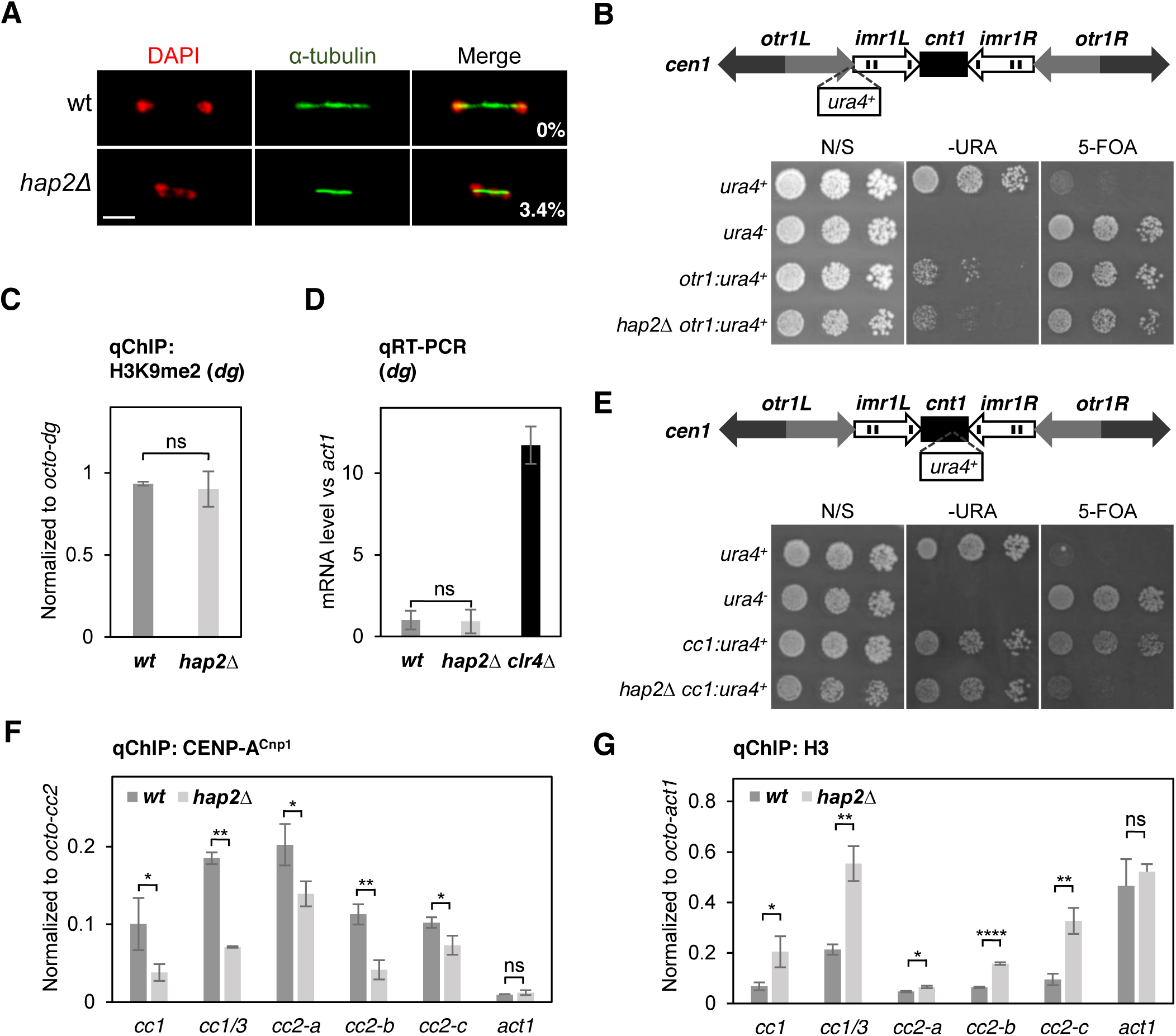
Hap2 is required to maintain CENP-A^Cnp1^ chromatin but not the adjacent pericentric heterochromatin. (A) Frequency of lagging chromosomes in anaphase cells. Representative images of wild-type and *hap2*Δ cells stained with DAPI (red) and anti-tubulin (green). Percentage of anaphase cells (n=100) displaying lagging chromosomes is indicated. Scale bar, 5 μm. (B) Top: Diagram indicating position *otr1:ura4^+^* marker gene inserted in outer repeat heterochromatin of *cen1*. Bottom: Growth assay for central domain silencing at *otr1:ura4^+^*. 5-fold serial dilution of cell cultures were spotted on non-selective (N/S), selective (−URA) or counter-selective (FOA) plates as indicated and incubated at 30°C for 3 days. (C) qChIP for H3K9me2 at outer repeat heterochromatin (*dg*). Error bars indicate mean ± SD (n = 3). (D) qRT-PCR of pericentric *dg* repeats performed on total RNA extracted from indicated strains. Transcript levels are shown relative to *act1^+^* and normalized to wild-type (n = 3). (E) Top: Diagram indicating position *cc1:ura4^+^* marker gene within the central CENP-A^Cnp1^ domain *cen1 (cnt1)* relative to the outer repeat (*otr*) and innermost repeats (*imr*). Bottom: Growth assay for central domain silencing at *cen1:ura4^+^* as in B. (F and G) qChIP for CENP-A^Cnp1^ (F) and histone H3 (G) at five locations within endogenous centromeres (*cc1*, cc1/3, *cc2-a*, *cc2-b* and *cc2-c*), outer repeat heterochromatin (*dg*) and non-centromere locus (*act1^+^*). Error bars indicate mean ± SD (n = 3). Significance of the differences observed between wild-type and *hap2*Δ was evaluated using Student’s t-test in C, D, F and G; * *p* < 0.05, ** *p* < 0.005; *** *p* < 0.005 and n.s., not significant.

Central CENP-A^Cnp1^ domain chromatin is also transcriptionally repressive (Allshire et al., 1994, 1995). Defects in CENP-A^Cnp1^ chromatin assembly alleviates this transcriptional silencing (Jin et al., 2002; Partridge et al., 2000; Pidoux et al., 2003). To test if loss of Hap2 affects CENP-A^Cnp1^-mediated silencing we examined silencing of *ura4^+^* embedded in central CENP-A^Cnp1^ chromatin at *cen1* (*cc1:ura4^+^*). Reduced silencing in *hap2*Δ cells, indicated by reduced growth on counter-selective FOA plates, suggested a defect in CENP-A^Cnp1^ chromatin integrity (Figures 3E and S3B). qChIP analysis detected significantly lower levels of CENP-A^Cnp1^ and a reciprocal increase in H3 levels across the central domain of centromeres in *hap2*Δ relative to wild-type cells (Figure 3F and G), consistent with the silencing defect. Importantly, loss of Hap2 does not affect the total cellular levels of GFP-CENP-A^Cnp1^ or the expression of genes encoding proteins that are known to regulate CENP-A^Cnp1^ loading at centromeres (Figure S3C and D). We conclude that loss of Hap2 specifically affects silencing through loss of CENP-A^Cnp1^ and gain of H3 within the central domain of centromeres, thus Hap2 is required to maintain CENP-A^Cnp1^ chromatin integrity at endogenous centromeres.

### Hap2 is required for the *de novo* establishment of CENP-A^Cnp1^ chromatin

Many factors are known to assist CENP-A maintenance but little is known about the factors required for the *de novo* establishment of CENP-A chromatin. *De novo* establishment of functional centromeres can occur following the introduction of naked centromere DNA into cells (Catania et al., 2015; Nakano et al., 2003). We first examined whether Hap2 and other subunits of Ino80 complex are required for the *de novo* establishment of centromeres on the pHcc2 minichromosome (Figure 4A). *hap2*Δ cells exhibited a complete failure to establish functional centromeres, similar to *clr4*Δ cells that lack heterochromatin, whilst *iec1*Δ, *ies2*Δ and *ies4*Δ were competent in establishing functional centromeres on pHcc2 (Figure 4B and S4A). In *S. pombe*, *de novo* CENP-A^Cnp1^ chromatin establishment on circular plasmid-based minichromosomes requires a block of heterochromatin in close proximity to central domain DNA (Folco et al., 2008; Kagansky et al., 2009). Loss of centromere establishment could result from a failure to establish CENP-A^Cnp1^ chromatin, and/or adjacent heterochromatin, on the minichromosome. To distinguish between these possibilities, CENP-A^Cnp1^ levels on the plasmid-borne central domain cc2 DNA were analysed. All strains used have 6 kb of *cen2* central domain DNA replaced with 5.5 kb of *cen1* central domain DNA at endogenous centromeres (*cc2Δ::cc1*) so that the plasmid-borne cc2 is the only copy of this element. qChIP revealed that CENP-A^Cnp1^ was not assembled over cc2 carried by pHcc2 in *hap2*Δ cells (Figure 4C). Reciprocally, a high level of H3 was detected across cc2 of pHcc2 in the absence of CENP-A^Cnp1^ assembly in *hap2*Δ cells (Figure 4D). qChIP for H3K9me2 demonstrated that loss of Hap2 did not affect the establishment of heterochromatin on the plasmid-borne K”/dg repeat (Figure 4E). Interestingly, high levels of H3K9me2 were detected within the central domain of the pHcc2 minichromosome in *hap2*Δ but not wild-type cells (Figure 4E). This suggests that in the absence of CENP-A^Cnp1^ chromatin establishment in *hap2*Δ cells heterochromatin may spread from the outer K”/dg repeat into the plasmid-borne central cc2 domain. To test if Hap2 is required for the *de novo* establishment of CENP-A^Cnp1^ chromatin independently from the requirement for adjacent heterochromatin, the plasmid pcc2 which carries 8.5 kb of *cen2* central domain DNA, but no heterochromatin forming outer repeat sequences, was transformed into cells expressing additional GFP-CENP-A^Cnp1^ which allows CENP-A^Cnp1^ chromatin assembly (Catania et al. 2015; Figure 4F, Top). In contrast to wild-type, *hap2*Δ cells did not assemble high levels of CENP-A^Cnp1^ over the central domain of the pcc2 minichromosome (Figure 4F, Bottom). This effect was not due to altered CENP-A^Cnp1^ protein levels in *hap2*Δ compared to wild-type cells (Figure S4B). We conclude that Hap2 is critical for the *de novo* establishment of CENP-A^Cnp1^ chromatin on naïve central domain centromere DNA.

**Figure 4.**
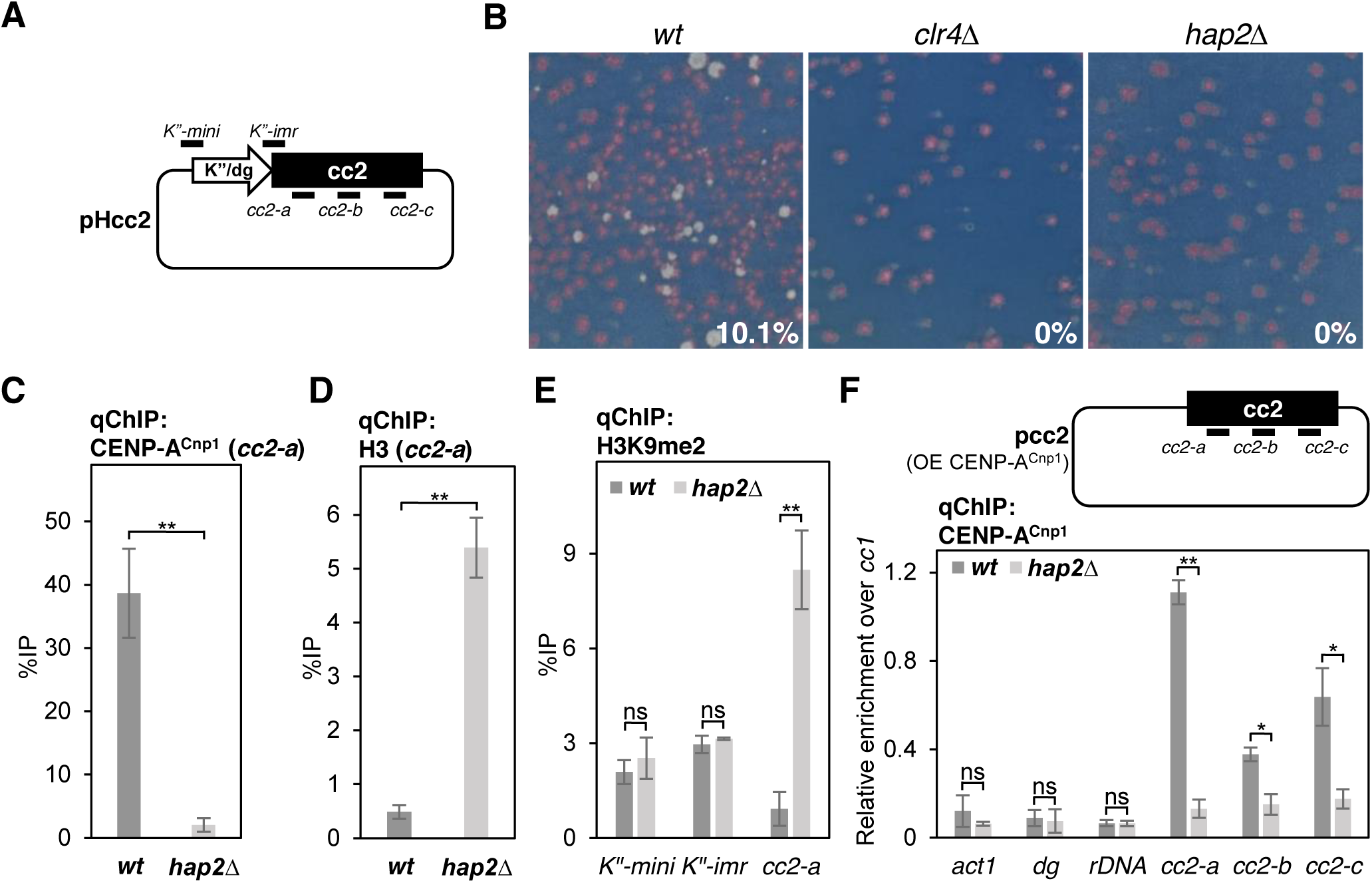
Hap2 is required for the *de novo* establishment of CENP-A^Cnp1^ chromatin. (A) Schematic representation of pHcc2 minichromosome containing 5.5 kb K”/dg repeat adjacent to 6.8 kb central domain 2 DNA. Position of the primer pair at the edge of K”/dg repeat (*K”-mini* and *K”-imr*) and within central domain 2 DNA (*cc2-a*, *cc2-b* and *cc2-c*) are indicated. (B) *S. pombe* transformants containing pHcc2 minichromosome plasmids were replica-plated to low adenine non-selective plates: colonies retaining pHcc2 minichromosome plasmid have established centromere function are white/pale-pink, those that lose it are red. Centromere establishment frequency with representative plate showing colonies with established centromere function and those that have failed to establish centromere function in wild-type (n = 515), *hap2*Δ (n = 390) and strains lacking heterochromatin *clr4*Δ (n = 870). (C and D) qChIP for CENP-A^Cnp1^ and histone H3 at plasmid-borne central domain 2 DNA (*cc2-a*) on pHCC2 minichromosome. Error bars indicate mean ± SD (n = 3). (E) qChIP for H3K9me2 at plasmid-borne K”/dg repeat (*K”-mini* and *K”-imr*) and plasmid-borne central domain 2 DNA (*cc2-a*) on pHcc2 minichromosome. Error bars indicate mean ± SD (n = 3). (F) Top: Schematic representation of pcc2 minichromosome without K”/dg repeat transformed in cells overexpressing CENP-A^Cnp1^. OE, overexpression. Bottom: qChIP for CENP-A^Cnp1^ at non-centromere locus (*act1^+^*), outer repeat heterochromatin (*dg*) and ribosomal DNA (*rDNA*) and three locations within plasmid-borne central core 2 DNA (*cc2-a*, *cc2-b* and *cc2-c*) on pcc2 minichromosome relative to CENP-A^Cnp1^ levels at endogenous centromeres (*cc1/3*). Error bars indicate mean ± SD (n = 3). Significance of the differences observed between wild-type and *hap2*Δ was evaluated using Student’s t-test in C, D, E and F; * *p* < 0.05, ** *p* < 0.005; *** *p* < 0.005 and n.s., not significant.

### Hap2 promotes histone turnover in genomic regions prone to CENP-A^Cnp1^ assembly

Central domain DNA inserted at a non-centromeric location on the arm of a chromosome, such as the *ura4* locus on chromosome 3 (*ura4:cc2*), remains assembled in H3 nucleosomes and exhibits a high rate of histone H3 turnover (Shukla et al., 2018). The inherent instability of H3 nucleosomes assembled on this ectopic *ura4:cc2* centromere DNA has been proposed to aid the incorporation of CENP-A^Cnp1^ when available (Shukla et al., 2018). In *S. cerevisiae*, Ino80 associates with promoter-associated nucleosome depleted regions and transcription start sites (TSS) where it mediates the turnover of +1 nucleosomes (Yen et al., 2013). The requirement for Hap2 in *de novo* CENP-A^Cnp1^ assembly on central domain DNA may result from defective H3 nucleosome turnover on these centromeric sequences when assembled in H3 chromatin alone. We therefore used <u>R</u>ecombination-Induced <u>T</u>ag <u>E</u>xchange (RITE; Verzijlbergen et al., 2010; Shukla et al., 2018) to measure replication-independent H3 turnover in G2-arrested wild-type and *hap2*Δ cells on ectopic *ura4:cc2,* heterochromatic repeats and highly-transcribed genes. This H3.2-HA→T7 tag swap was induced in *cdc25-22*/G2-arrested cells and the incorporation of new histone H3.2-T7 was monitored (Figure 5A). Importantly, the H3.2-HA→T7 tag swap efficiency was unaffected by *hap2*Δ relative to wild-type cells (Figure S5A). Compared to wild-type cells, a significant decrease in the level of H3 turnover was evident on ectopic *ura4:cc2* centromere DNA in *hap2*Δ cells (Figure 5B). Similarly, H3 turnover within endogenous *cen1-cc1* assembled in CENP-A^Cnp1^ chromatin was also reduced in *hap2*Δ cells. In contrast, H3 turnover remained unchanged within heterochromatic outer repeats (*dg*) and over highly-transcribed genes (*act1^+^* and *spd1^+^*). Ino80 may mediate nucleosome turnover through eviction of H2A.Z (Papamichos-Chronakis et al., 2011). However, the levels of H2A.Z^Pht1^ associated with endogenous *cc1* and ectopic *ura4:cc2* were unaffected by loss of Hap2 (Figure S5B). We conclude that Hap2 is required to ensure H3 nucleosome instability on centromeric sequences by mediating a high frequency of H3 turnover which may consequently allow the incorporation of CENP-A^Cnp1^ in place of H3.

**Figure 5.**
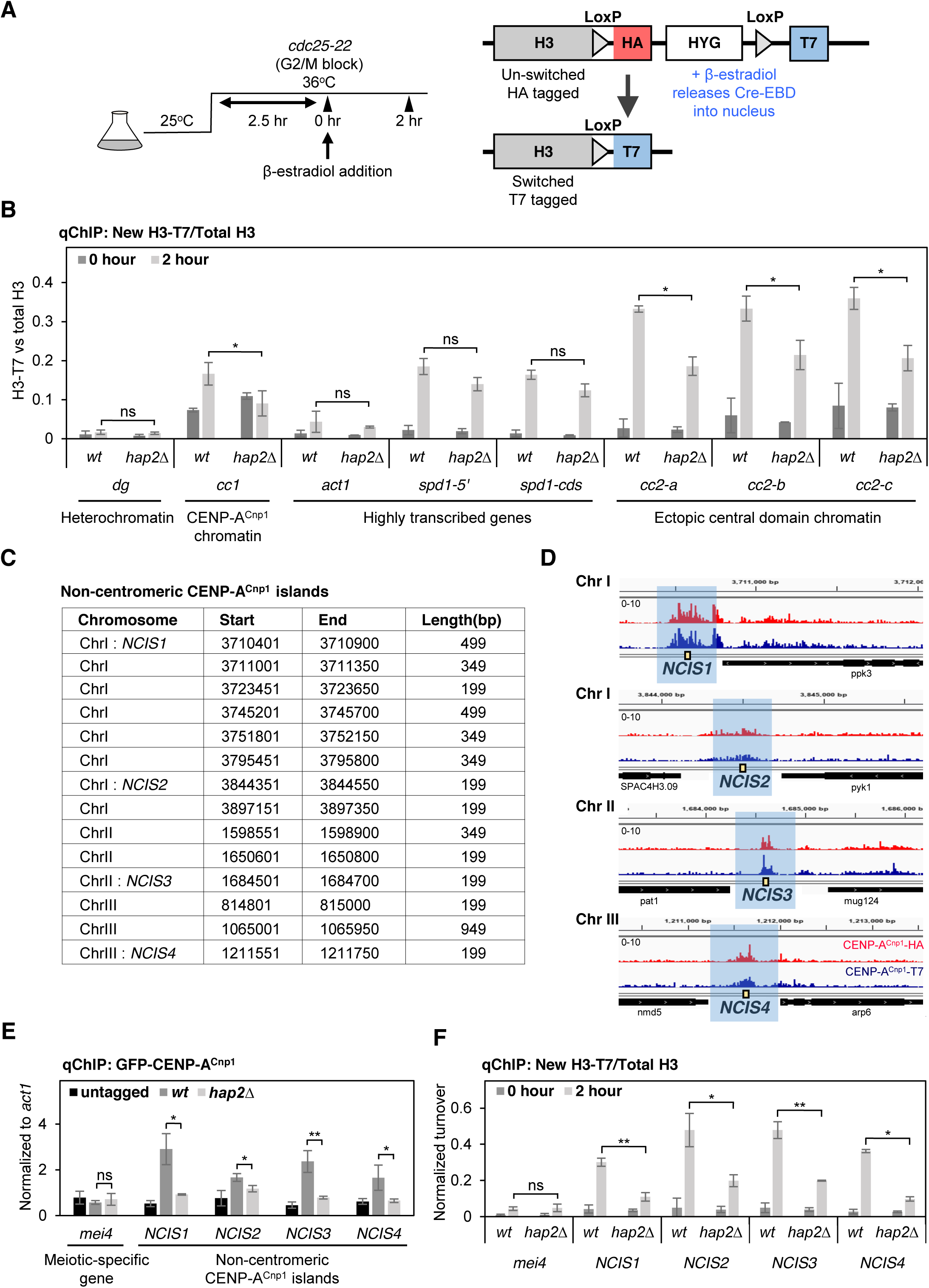
Hap2 is required for high histone H3 turnover on central domain centromere DNA and islands of non-centromeric CENP-A incorporation. (A) Experimental setup to assess replication-independent H3 turnover. The c*dc25-22* temperature sensitive mutation was used to block wild-type and *hap2*Δ cells in G2, after 2.5 hours at 36°C, the HA tag on H3 was swapped for the T7 tag by β-estradiol induced Cre/loxP mediated recombination. Samples were collected at 0 and 2 hours and analyzed by qChIP. (B) qChIP analysis of new H3.2-T7 incorporated in *cdc25-22*/G2 arrested cells at pericentromeric outer repeat heterochromatin (*dg)*, the central CENP-A^Cnp1^ domain of cen1 (*cc1)*, highly transcribed genes (*act1^+^* and *spd1^+^*) and three locations within ectopically located *ura4:cc2* lacking CENP-A^Cnp1^ (*cc2-a*, *cc2-b* and *cc2-c*). y axis: H3.2-T7 %IP values normalized to the respective total H3 %IP values represents normalized turnover for each sample. Error bars indicate mean ± SD (n = 3). (C) ChIP-seq analysis of HA and T7 tagged CENP-A^Cnp1^ reveals consistent <u>N</u>on-centromeric <u>C</u>ENP-A^Cnp1^ <u>IS</u>lands (NCIS) of incorporation above background at 14 locations in the genome. (D) Four CENP-A^Cnp1^ islands are shown: *NCIS1*, *NCIS2*, *NCIS3* and *NCIS4* (data from C). (E) qChIP for GFP-CENP-A^Cnp1^ at non-centromeric CENP-A^Cnp1^ islands using indicated primer pairs (yellow boxes in D for *NCIS1*, *2*, *3* and *4*) and meiotic-specific gene (*mei4^+^*). Error bars indicate mean ± SD (n = 3). (F) qChIP analysis of new H3.2-T7 incorporation (turnover as in B) in *cdc25-22*/G2 arrested cells at *NCIS1*, *2*, *3* and *4*. Error bars indicate mean ± SD (n = 3). Error bars indicate mean ± SD (n = 3). Significance of the differences observed between wild-type and *hap2*Δ was evaluated using Student’s t-test in B, E and F; * *p* < 0.05, ** *p* < 0.005; *** *p* < 0.005 and n.s., not significant.

Overexpression of CENP-A^Cnp1^ results in low levels of promiscuous CENP-A^Cnp1^ incorporation at non-centromeric locations (Castillo et al., 2013; Choi et al., 2012; Gonzalez et al., 2014). Fission yeast CENP-A^Cnp1^ expression is known to increase prior to that of canonical histones in advance of replication. Consequently, even without overexpression, in early S phase this natural excess of CENP-A^Cnp1^ results in low levels of newly synthesized CENP-A^Cnp1^ being incorporated across many gene bodies (Shukla et al., 2018). ChIP-Nexus (a modified Exo-ChIP-seq protocol; He et al., 2015) analysis allowed detection of non-centromeric genomic regions where islands of CENP-A^Cnp1^ are retained at low levels (Figure 5C and D). Hap2-GFP was found to associate with these <u>N</u>on-centromeric <u>C</u>ENP-A^Cnp1^ <u>IS</u>lands (NCIS), while loss of Hap2 significantly decreased CENP-A^Cnp1^ incorporation within these islands (Figure 5E and S5C). Interestingly, qChIP also revealed reduced histone H3 turnover within these non-centromeric CENP-A^Cnp1^ islands in *hap2*Δ cells whereas histone H3 turnover remained unchanged within the outer repeats, the *act1^+^* gene or the repressed meiosis-specific *mei4^+^* gene (Figure 5F). We conclude that Hap2 also promotes a high rate of H3 turnover at several non-centromeric NCIS loci prone to low level CENP-A^Cnp1^ incorporation. This observation reinforces the involvement of Hap2-Ino80C in mediating histone H3 turnover and the exchange of histone H3 for CENP-A^Cnp1^-containing nucleosomes.

### Hap2 facilitates transcription of central domain chromatin

The central CENP-A^Cnp1^ domain of *S. pombe* centromeres is transcribed from many TSS and the resulting RNAs are short-lived (Catania et al., 2015; Choi et al., 2011; Sadeghi et al., 2014). Transcription can provide the opportunity for histone exchange/remodeling of resident nucleosomes (Venkatesh and Workman, 2015), it is therefore feasible that transcription-coupled processes also promote the exchange of H3 for CENP-A^Cnp1^ in chromatin assembled on centromere DNA. Relatively high levels of RNAPII are detected on central domain DNA when assembled in H3 chromatin at ectopic *ura4:cc2* or on the pcc2 minichromosome, yet only low levels of RNAPII are detectable within endogenous centromeric central domain CENP-A^Cnp1^ chromatin (Catania et al., 2015; Shukla et al., 2018). Since CENP-A^Cnp1^ is primarily deposited during G2, we investigated whether Hap2 affects transcription from ectopic *ura4:cc2* in mid-G2 cells from *cdc25-22* synchronized cultures (Figure 6A). qChIP was employed to measure the levels of total RNAPII, initiating RNAPIIS5P and elongating RNAPIIS2P levels across *ura4:cc2*. RNAPIIS5P levels were significantly lower across this ectopic central domain DNA in *hap2*Δ compared to wild-type cells, but no obvious difference was detected within the CENP-A^Cnp1^ chromatin regions of endogenous centromeres (Figure 6B). These data suggest that Hap2 promotes efficient transcriptional initiation from central domain DNA assembled in H3 chromatin (*ura4:cc2*). In contrast, the levels of total and elongating RNAPIIS2P associated with ectopic central domain DNA remained unchanged in *hap2*Δ relative to wild-type cells (Figure S6A and B). Consistent with reduced transcriptional initiation, lower levels of central domain transcripts were produced from ectopic *ura4:cc2* in *hap2*Δ cells (Figure 6C). Transcript levels from outer repeats and *act1^+^* were unaffected. Thus, Hap2 is required for efficient transcriptional initiation and transcript production from ectopic central domain DNA assembled in H3 chromatin. The fact that no decrease in elongating RNAPIIS2P was detected across this ectopic central domain in *hap2*Δ cells suggests that RNAPII remains associated with the central domain template for longer when Hap2 is absent.

**Figure 6.**
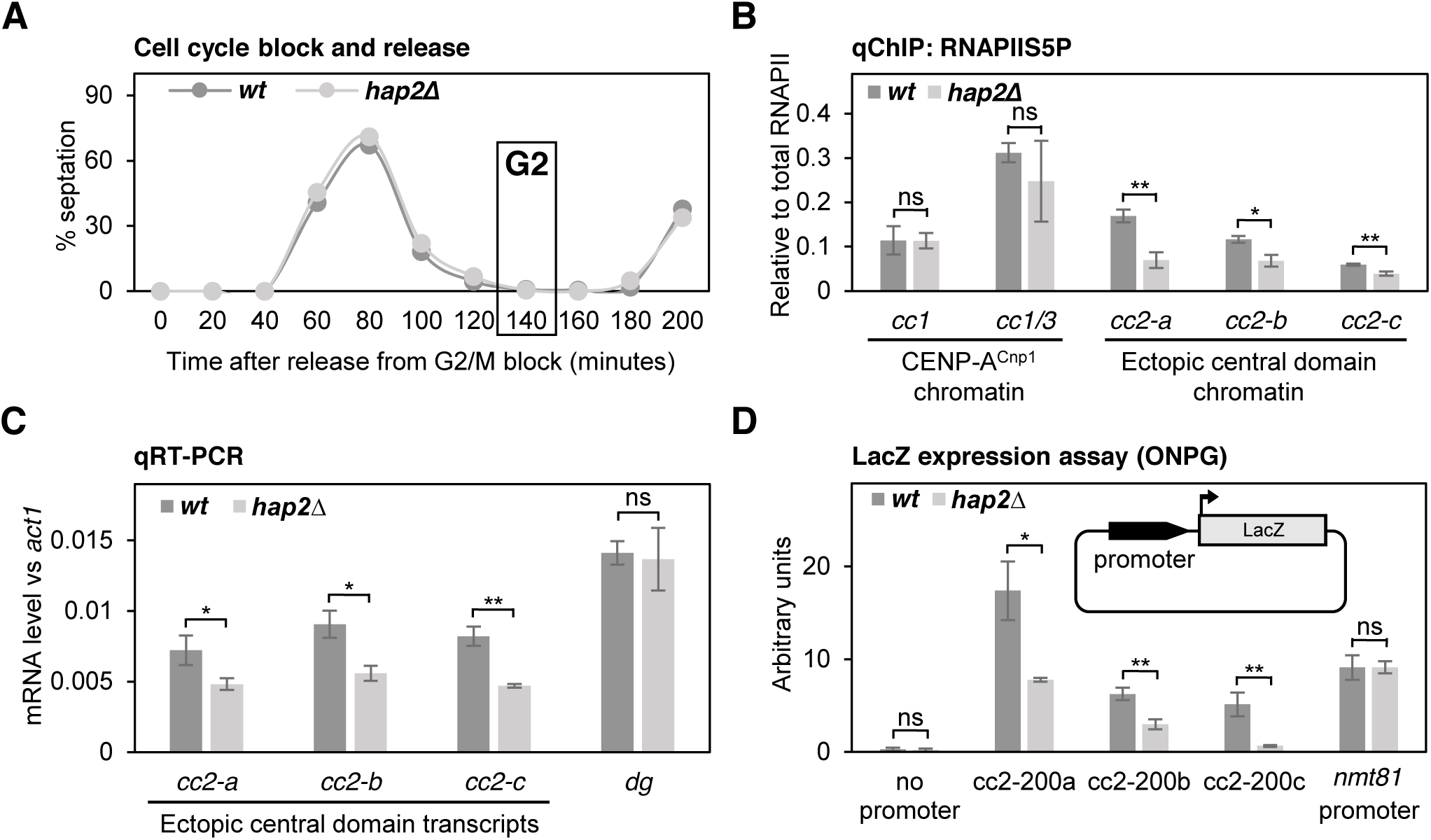
Transcription from centromeric DNA is reduced in the absence of Hap2. (A) *cdc25-22* synchronized cell populations were used to assess levels of RNAPIIS5P and central core transcripts during G2 in B and C. The septation index for wild-type and *hap2*Δ were measured and cells were collected in G2 (Time point: 140) for further analysis in B and C. (B) qChIP for initiating RNAPIIS5P at two locations within endogenous centromeres (*cc1, cc1/3*) and three locations within ectopically located *ura4:cc2* lacking CENP-A^Cnp1^ (*cc2-a*, *cc2-b* and *cc2-c*) in wild-type and *hap2*Δ. Error bars indicate mean ± SD (n = 3). (C) qRT-PCR to quantify transcripts from three locations within ectopically located *ura4:cc2* lacking CENP-A^Cnp1^ (*cc2-a*, *cc2-b* and *cc2-c*) and from pericentromeric outer repeat heterochromatin (*dg)* on total RNA extracted from in wild-type and *hap2*Δ. Transcript levels are shown relative to *act1^+^,* error bars indicate mean ± SD (n = 3). (D) Measurement of LacZ expression driven by three 200bp *cc2* fragments with promoter activity (cc2-200a, cc2-200b and cc2-200c), the nmt81 promoter and no promoter following transformation of plasmids into wild-type and *hap2*Δ using the ONPG substrate/absorbance at 420 nm, error bars indicate mean ± SD (n = 3). Inset: Diagram of plasmids with different 200 bp fragments from central domain region of *cen2* (cc2) placed upstream the LacZ reporter. Significance of the differences observed between wild-type and *hap2*Δ was evaluated using Student’s t-test in B-D; * *p* < 0.05, ** *p* < 0.005; *** *p* < 0.005 and n.s., not significant.

Regions upstream of TSS within the central domain of *cen2* exhibit promoter activity (Catania et al., 2015). To determine if Hap2 affects the transcription from these central domain promoters, the production of β-galactosidase was assessed when 200 bp promoter-containing cc2 fragments were placed upstream of lacZ (Figure 6D Inset). Three central domain promoters exhibited significantly lower promoter activity in *hap2*Δ compared to wild-type cells, whereas the control *nmt81* promoter was unaffected (Figure 6D; S6C and D). We conclude that Hap2 is required for efficient transcription from central domain promoters and that this facilitates transcription-coupled histone exchange, thereby providing an opportunity for replacement of H3 with CENP-A^Cnp1^ to establish CENP-A^Cnp1^ chromatin domains and assemble functional kinetochores.

## DISCUSSION

The mechanisms that contribute to the maintenance and, especially, the establishment, of CENP-A chromatin remain poorly understood. To gain insight into how CENP-A chromatin is established on naïve centromere DNA, we applied proteomics to identify proteins associated with fission yeast CENP-A^Cnp1^ chromatin. In addition to kinetochore proteins, all Ino80C subunits, including the small auxiliary subunit, Hap2, were found to be significantly enriched in solubilized CENP-A^Cnp1^ chromatin. Hap2 was found to promote CENP-A^Cnp1^ chromatin integrity at centromeres and to be required for the *de novo* establishment of CENP-A^Cnp1^ chromatin on introduced naïve centromere DNA. The requirement for Hap2 in ensuring high histone H3 turnover on endogenous centromere DNA, ectopically-located centromere DNA, and non-centromeric CENP-A^Cnp1^ islands indicates that Hap2-Ino80C drives H3 nucleosome turnover at these locations. The loss of CENP-A^Cnp1^ incorporation from NCIS islands in the absence of Hap2 underscores the role for Hap2-Ino80C-mediated H3 turnover in stimulating CENP-A^Cnp1^ incorporation. Strikingly, Hap2 is required for efficient transcription specifically from central domain promoters. We propose a mechanism in which Hap2-Ino80C drives the inherent instability of H3 nucleosomes on centromeric DNA via transcription-coupled nucleosome turnover, providing the opportunity for incorporation of CENP-A^Cnp1^ in place of histone H3, when CENP-A^Cnp1^ is available (Figure 7).

**Figure 7.**
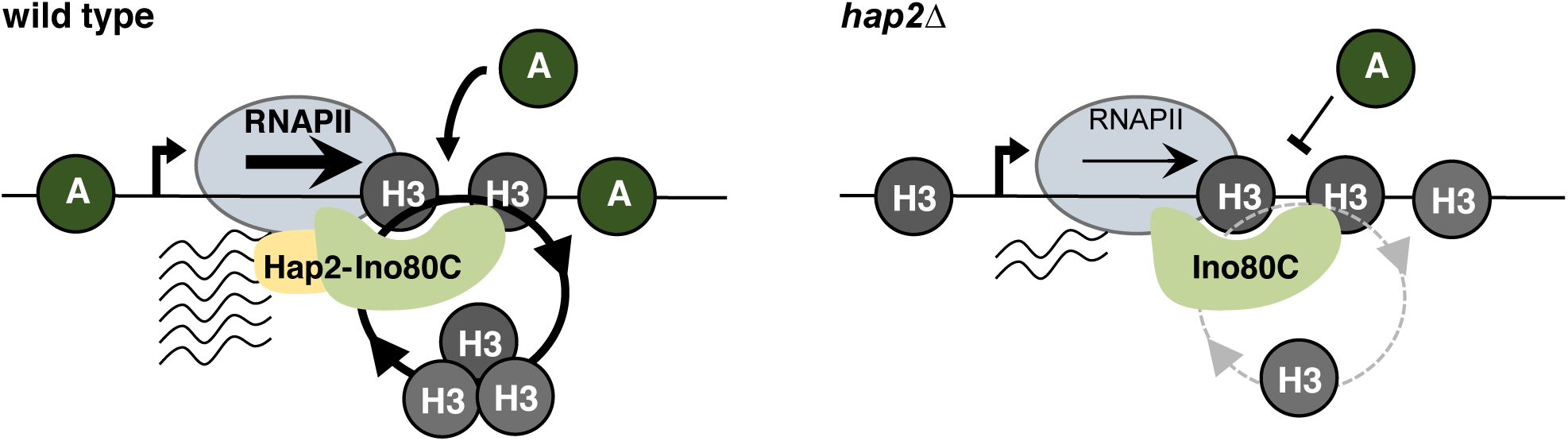
Hap2-Ino80 mediated transcription facilitates *de novo* establishment of CENP-A^Cnp1^ chromatin. Model for the establishment of CENP-A^Cnp1^ chromatin in *S. pombe*. Left: Hap2 is required for transcription initiation events to maintain high H3 turnover thus renders H3 nucleosomes unstable on centromeric sequences allowing its replacement with CENP-A^Cnp1^ nucleosomes. Right: Cells lacking Hap2 have lower transcription initiation events that stabilize the H3 nucleosomes on centromeric sequences thus lowering its propensity to establish CENP-A^Cnp1^ chromatin.

Studies in a variety of species have shown that Ino80C influences transcription (Cai et al., 2007; Hogan et al., 2010; Klopf et al., 2009; Neuman et al., 2014; Steger et al., 2003). The non-canonical human Yin Yang 1 (YY1) transcription factor is known to associate with Ino80 and facilitate both transcriptional activation and repression (Cai et al., 2007; Lee et al., 1995; Yao et al., 2001). It is also known that Ino80 suppresses anti-sense and other non-coding transcription (Alcid and Tsukiyama, 2014; Marquardt et al., 2014). The non-coding transcription of centromeric DNA by RNAPII has been implicated in CENP-A deposition in several systems (Bobkov et al., 2018; Chan et al., 2012; Chueh et al., 2009; Grenfell et al., 2016; McNulty et al., 2017; Rošić et al., 2014). However, both reduced and increased transcription of centromere DNA appears to be incompatible with CENP-A chromatin integrity and centromere function (Bergmann et al., 2012; Hill and Bloom, 1987; Nakano et al., 2008; Ohkuni and Kitagawa, 2011). Such observations suggest that an appropriate level and type of programmed transcription may be required to promote CENP-A assembly on centromere DNA.

Each fission yeast centromere contains a central domain of approximately 10 kb assembled in CENP-A^Cnp1^ rather than H3 nucleosomes. Multiple transcriptional start sites are detected on both strands when central domain is assembled in H3, rather than CENP-A^Cnp1^, chromatin at an ectopic locus (Choi et al. 2011; Catania et al. 2015), indicating that these regions are pervasively transcribed. Distinct Ino80C modules undertake particular tasks such as nucleosome binding, sliding and ATPase activity (Zhou et al., 2016). The specific impact of Hap2 on Ino80C remains to be determined. However, a major activity of Ino80C is to slide nucleosomes relative to sizable lengths of flanking unoccupied DNA (Udugama et al., 2011). Thus, the generation of nucleosome free regions that facilitate RNAPII recruitment, and consequently transcription, are an intrinsic facet of Ino80C function. Consequently, the loss of Hap2-Ino80C is expected to occlude central domain promoters with nucleosomes and result in reduced transcriptional initiation.

Since RNAPII recruitment to central domain chromatin during G2 phase of the cell cycle is coincident with H3 eviction and CENP-A^Cnp1^ incorporation (Shukla et al., 2018), it is likely that these events are somehow coupled. The conserved heptad repeat composing the C-terminal domain (CTD) of RNAPII undergoes S5 phosphorylation (S5P) at promoters upon transcription initiation and S2 phosphorylation (S2P) in coding regions during transcriptional elongation (Komarnitsky et al., 2000). RNAPII stalls when it encounters obstacles such as DNA damage or natural barriers (Poli et al., 2017). Despite relatively high levels of RNAPII being detected on H3-assembled ectopic central domain, meagre levels of transcripts are produced, consistent with transcriptional stalling (Catania et al., 2015; Choi et al., 2011). Analyses in *S. cerevisiae* show that Ino80 is required to release stalled RNAPII from chromatin and enable its proteasomal degradation (Lafon et al., 2015). Previously, we showed that mutants (*ubp3*Δ, *tfs1*Δ) expected to increase the levels of stalled RNAPIIS2P on central domain chromatin, promote CENP-A incorporation (Catania et al., 2015). Interestingly, cells lacking Hap2 display lower initiating RNAPIIS5P over ectopic H3-assembled central domain chromatin in G2 but the levels of total RNAPII and elongating RNAPII-S2P are unaltered (Fig. 6B, S6A and B). This indicates that, although lower levels of initiation and transcription take place within ectopic central domain chromatin in the absence of Hap2, elongating RNAPII is retained across the domain. We interpret these observations to indicate that Hap2-Ino80C is required to release elongating RNAPII that becomes trapped or stalled in central domain chromatin. We suggest that resident H3 nucleosomes are evicted by Hap2-Ino80C as part of the process involved in removing this stalled RNAPII. Thus, centromere DNA may be programmed to be pervasively transcribed and stall RNAPII over a relatively large 10 kb region in order to recruit Hap2-Ino80C and trigger H3 turnover throughout this extensive domain, thereby providing the opportunity for CENP-A incorporation. Stalling may result from collisions between converging RNAPII complexes or other obstacles such as the replication Origin Recognition Complex (ORC) which is bound to the many AT-rich tracts present within central domain regions (Hayashi et al., 2007). Alternatively, elongating RNAPII on central domain DNA may be somehow earmarked for removal using cues analogous to those that allow Ino80 to suppresses anti-sense and cryptic unstable transcripts (Alcid and Tsukiyama, 2014). Thus, pervasive transcription across an extensive region such as the central domain may itself provoke Ino80C recruitment.

Histone turnover rates are generally higher for nucleosomes close to promoters (Dion et al., 2007; Jin and Felsenfeld, 2007; Zhang et al., 2005). The first +1 nucleosomes downstream of many eukaryotic promoters contain the H2A.Z variant (rather than canonical H2A) to aid transcription. These +1 nucleosomes exhibit high levels of turnover and Ino80C has been reported to mediate the exchange of H2A.Z for canonical H2A in such nucleosomes (Papamichos-Chronakis et al., 2011; Watanabe et al., 2013; Yen et al., 2013). However, H2A.Z turnover on transcribed genes has been shown to be not reliant on Ino80 activity (Tramantano et al., 2016). Moreover, Ino80 is also known to reduce transcription from some promoters, independently of its role in removing H2A.Z (Barbaric et al., 2007). The fact that no difference in the levels of H2A.Z^Pht1^ was detected across the ectopic central domain in *hap2*Δ cells relative to wild-type (Figure S5B) suggests that Hap2-Ino80C does not mediate H2A.Z^Pht1^ eviction from central domain chromatin and that it must affect some other aspect of centromere promoter function, perhaps through its known nucleosome sliding activity. Thus, Ino80C may be required to slide nucleosomes away from central domain promoters so that they are efficiently transcribed and the resulting transcriptional properties of this domain mediate H3 nucleosome turnover.

Previously we showed that CENP-A^Cnp1^ competes with histone H3 for incorporation into centromeric chromatin and CENP-A^Cnp1^ incorporation is promiscuous, replacing H3 when the opportunity arises (Castillo et al., 2007; Choi et al., 2011, 2012). Thus, high histone H3 nucleosome turnover, or low nucleosome occupancy, appears to be an underlying property of many sequences that are prone to CENP-A^Cnp1^ assembly in fission yeast. As transcription appears to be widespread at centromeres in other organisms, high turnover of resident H3 nucleosomes, stimulated by Ino80C, may be a conserved attribute of centromeric DNA that stimulates CENP-A deposition.

## MATERIALS AND METHODS

### Cell growth and manipulation

Standard genetic and molecular techniques were followed. Fission yeast methods were as described (Moreno et al., 1991). Fission yeast strains are listed in Table S3. YES (Yeast Extract with Supplements) was used as a rich medium or PMG (Pombe Minimal Glutamate) for growth of cells in liquid cultures. 4X-YES was used for experiments where higher cell numbers were required. H3.2 RITE strains containing HA/T7 RITE cassettes were as described (Shukla et al., 2018; Verzijlbergen et al., 2010). Plasmids used in this study are listed in Table S4.

### Native immunoaffinity purification

Cell were grown in 4X-YES and 2-4×10^10^ cells were used for immunoprecipitation. Cells were harvested by centrifugation at 3500×g, washed twice with water and flash frozen in liquid nitrogen. Frozen cell pellets were ground using Retsch MM400 mill. The grindate was resuspended in lyisis buffer (10 mM Tris pH7.4, 5 mM CaCl2, 5 mM MgCl2, 50 mM NaCl, 0.1% IGEPAL-CA630 and supplemented with protease inhibitor (P8215, Sigma) and 2 mM PMSF and thawed for 30 minutes on ice. Chromatin was solubilized by incubation with 4-20 units of Micrococcal nuclease (N3755, Sigma) for 10 minutes at 37° C. MNase digestion was stopped by adding EGTA to 20 mM and lysates were rotated at 4° C for 1 hour to ensure chromatin solubilization. Supernatant was separated from lysates by centrifugation at 20,000×g for 10 minutes and used for immunoprecipitation using 20 µg of anti-GFP antibody (11814460001, Roche) (used for GFP-CENP-A^Cnp1^ and Hap2-GFP IP) or 20 µg of anti-HA antibody (ab9110, Abcam) (used for Ino80-3HA IP) coupled to 50 µl of Protein G Dynabeads (Life Technologies) using dimethyl pimelimidate (DMP; 21666, ThermoFisher Scientific). Bead-bound affinity-selected proteins were washed three times with lysis buffer. Elution was performed with 0.1% RapiGest SF (186001860, Waters) in 50mM Tris-HCl pH8 by incubating at 50° C for 10 minutes.

### Co-immunoprecipitation

The co-immunoprecipitation protocol is similar to native immunoaffinity purification described above with following modifications. Cell were grown in 4X-YES and 5×10^9^ cells were used per IP. Chromatin was solubilized by incubation with 100 units of Micrococcal nuclease (Sigma, N3755) for 20 minutes at 37° C. Immunoprecipitation was performed using 10 µg of anti-GFP antibody (Roche, 11814460001) (used for Hap2-GFP IP) or 10 µg of anti-HA antibody (Abcam, ab9110) (used for Ino80-3HA IP) coupled to 25 µl of Protein G Dynabeads (Life Technologies) using DMP. Bead-bound affinity-selected proteins were washed five times with lysis buffer for 5 minute and eluted with LDS loading buffer (ThermoFisher Scientific, 84788) by incubating at 75° C for 10 minutes. Western blotting detection was performed using anti-GFP (ThermoFisher Scientific, A11122) or anti-HA antibody (Abcam, ab9110) and HRP conjugated anti-Rabbit IgG HRP (Sigma, A6154)).

### Mass spectrometry and analysis

Proteins were digested with trypsin into peptides and analyzed by mass spectrometry. Proteins were denatured by adding final 25mM DTT to the immunoprecipitated proteins and incubated at 95° C for 5 minutes. Samples were cooled at room temperature and 200 µl of 8M Urea in 100mM Tris-HCl pH8 was added and passed through Vivacon 500 column (VN01H21, Sartorius Vivacon 500, 30,000 MWCO Hydrosart) by centrifugation at 14,000xg for 30 minutes. Cysteine residues were alkylated to prevent free sulfhydryls from reforming disulfide bonds by incubation with 100 µl of 0.05 M Iodoacetamide (Sigma) in 8 M urea for 20 minutes at 27° C in the dark and centrifuged at 14,000xg to remove the supernatant. Columns were then washed once with 100 µl of 8M urea and twice with 100 µl of 0.05 M ammonium bicarbonate (Sigma). Proteins were digested by addition of 100 µl of 0.05 M ABC containing 0.3 µg of trypsin (90057, Thermo Scientific) at 37° C for 16 hours. Columns were spun at 14,000xg to collect the digested peptides and washed again with 100 µl of 0.05 M ABC. Trypsinization was stopped by addition of 10 µl of 10% trifluoroacetic acid (TFA) to bring the pH to below 2.

C18 reverse-phase resin (66883-U, Sigma) was used to desalt peptide samples prior to LC-MS/MS. StageTips were packed tightly with 2 layers of C18 resin. The resin was conditioned with 30 µl of 100% methanol, washed with 30 µl of 80% acetonitrile to remove impurities and finally equilibrated by passing 30 µl of 0.1% TFA. The trypsinized peptide solution was passed through the stage tip by centrifugation at 2600 rpm for binding. Following stage tip desalting, samples were diluted with equal volume of 0.1% TFA and spun onto StageTips as described (Rappsilber et al., 2003). Peptides were eluted in 40 μL of 80% acetonitrile in 0.1% TFA and concentrated down to 2 μL by vacuum centrifugation (Concentrator 5301, Eppendorf, UK). The peptide sample was then prepared for LC-MS/MS analysis by diluting it to 5 μL by 0.1% TFA.

LC-MS-analyses were performed on an Orbitrap Fusion™ Lumos™ Tribrid™ Mass Spectrometer and on a Q Exactive (both from Thermo Fisher Scientific, UK) both coupled on-line, to an Ultimate 3000 RSLCnano Systems (Dionex, Thermo Fisher Scientific, UK). In both cases, peptides were separated on a 50 cm EASY-Spray column (Thermo Scientific, UK), which was assembled on an EASY-Spray source (Thermo Scientific, UK) and operated at 50° C. Mobile phase A consisted of 0.1% formic acid in LC-MS grade water and mobile phase B consisted of 80% acetonitrile and 0.1% formic acid. Peptides were loaded onto the column at a flow rate of 0.3 μL min^−1^ and eluted at a flow rate of 0.2 μL min^−1^ according to the following gradient: 2 to 40% mobile phase B in 150 min and then to 95% in 11 min. Mobile phase B was retained at 95% for 5 min and returned back to 2% a minute after until the end of the run (190 min).

For Fusion Lumos, FTMS spectra were recorded at 120,000 resolution (scan range 350-1500 m/z) with an ion target of 7.0×10^5^. MS2 was performed in the ion trap with ion target of 1.0×10^4^ and HCD fragmentation (Olsen et al., 2007) with normalized collision energy of 27. The isolation window in the quadrupole was 1.4 Thomson. Only ions with charge between 2 and 7 were selected for MS2. For Q Exactive, FTMS spectra were recorded at 70,000 resolution (scan range 350-1400 m/z) and the ten most intense peaks with charge ≥ 2 of the MS scan were selected with an isolation window of 2.0 Thomson for MS2 (filling 1.0×10^6^ ions for MS scan, 5.0×10^4^ ions for MS2, maximum fill time 60 ms, dynamic exclusion for 50 s).

The MaxQuant software platform (Cox and Mann, 2008) version 1.5.2.8 was used to process the raw files and search was conducted against *Schizosaccaromyces pombe* complete/reference proteome set of PomBase database (released in July, 2015), using the Andromeda search engine (Cox et al., 2011). For the first search, peptide tolerance was set to 20 ppm while for the main search peptide tolerance was set to 4.5 pm. Isotope mass tolerance was 2 ppm and maximum charge to 7. Digestion mode was set to specific with trypsin allowing maximum of two missed cleavages. Carbamidomethylation of cysteine was set as fixed modification. Oxidation of methionine, phosphorylation of serine, threonine and tyrosine and ubiquitination of lysine were set as variable modifications. Label-free quantitation analysis was performed by employing the MaxLFQ algorithm as described by Cox et al. 2014. Absolute protein quantification was performed as described in Schwanhüusser et al. 2011. Peptide and protein identifications were filtered to 1% FDR.

The Perseus software platform (Tyanova et al., 2016) version 1.5.5.1 was used to process LFQ intensities of the proteins generated by MaxQuant. The LFQ intensities were transformed to log2 scale and list was filtered for proteins with at least 2-3 valid values in any sample. Missing values were imputed from normal distribution for that IP.

### Cytology

Cells were fixed with 3.7% formaldehyde (Sigma) for 7 minutes at room temperature. Immunolocalization staining was performed as described (Shukla et al., 2018). The following antibodies were used at 1:100 dilution: Anti-CENP-A^Cnp1^ (sheep in-house), anti-GFP (ThermoFisher Scientific, A11122), anti-TAT1 (mouse in-house); Alexa 594 and 488 labelled secondary antibodies at 1:1000 dilution (Life Technologies). Images were acquired with Zeiss LSM 880 confocal microscope equipped with Airyscan superresolution imaging module, using a 100X/1.40 NA Plan-Apochromat Oil DIC M27 objective lens and processed using ZEN Black image acquisition and processing software (Zeiss MicroImaging).

### Establishment assay

Minichromosomes used in this study were transformed using sorbitol electroporation method (Suga and Hatakeyama, 2001). Transformants were plated on PMG-uracil-adenine plates and incubated at 32°C for 5-10 days until medium-sized colonies had grown. Colonies were replica-plated to PMG low adenine (10 µg/ml) plates to determine the frequency of establishment of centromere function. These indicator plates allow minichromosome loss (red colony) or retention (white/pale pink colony) to be determined. Minichromosome retention indicates that centromere function has been established and that minichromosomes segregate efficiently in mitosis. In the absence of centromere establishment, minichromosomes behave as episomes that are rapidly lost. Minichromosomes occasionally integrate giving a false positive white phenotype. To assess the frequency of such integration events and to confirm establishment of centromere segregation function, colonies giving the white/pale-pink phenotype upon replica plating were re-streaked to single colonies on low-adenine plates. Sectored colonies are indicative of segregation function with low levels of minichromosome loss, whereas pure white colonies are indicative of integration into endogenous chromosomes and the establishment frequency was adjusted accordingly.

### LacZ assay

LacZ assay was performed as described (Guarente, 1983). Plasmids containing LacZ with upstream nmt81 promoter, 200 bp sequences from centromere 2 or no promoter were used as described (Catania et al., 2015). Plasmids were transformed into wild-type and *hap2*Δ strains and grown on minimal medium.

### ChIP-qPCR

Cells were grown at appropriate temperature and medium. Cultures were fixed with 3.7% formaldehyde for 15 min at room temperature. ChIP was performed as previously described (Castillo et al., 2007). Cells were lysed by bead-beating (Biospec Products) and sonicated in a Bioruptor (Diagenode) (20 min, 30 s On and 30 s Off at ‘High’ (200 W). 10 µl anti-CENP-A^Cnp1^ (sheep in-house antiserum) and 25 µl Protein-G-Agarose beads (Roche) or 2 µl of anti-GFP (A11122, ThermoFisher Scientific), 2 µl of anti-H3 (ab1791, Abcam), 2 µl of anti-T7 (69522, Millipore), 2 µl of anti-myc (2276, Cell Signaling Technology), 1 µl of anti-H3K9me (m5.1.1), 2 µl of anti-RNAPII (ab817, Abcam), 2 µl of RNAPII S2P (Abcam, ab5095) and 2 µl of RNAPII S5P (61085, Actif motif) and 20 µl Protein-G-Dyna beads (Life Technologies) were used per ChIP. qPCR was performed using Light Cycler 480 SybrGreen Master Mix (Roche, 04887352001) and analysed using Light Cycler 480 Software 1.5 (Roche). Primers used in qPCR are listed in Supplementary Table S5.

### qRT-PCR

RNA was isolated with RNeasy mini kit (Qiagen) according to the manufacturer’s protocol. A total of 10 µg RNA isolated by RNeasy Mini kit (Qiagen) was treated with 1 µl of Turbo DNase (AM2238, Invitrogen) for 30 minutes at 37°C. A second digestion was followed by adding additional 1 µl of Turbo DNase for 30 minutes at 37°C. RNA was then cleaned up according to RNeasy Mini Protocol for RNA Cleanup (Qiagen) and dissolved in nuclease free water. For qRT-PCR analysis, first strand cDNA synthesis was performed using 100 ng of random hexamer (Invitrogen), 1 µg of DNA-free RNA template and 1 µl of Superscript III (Invitrogen) reverse transcriptase according to the manufacturer’s instruction. As a negative control (-RT), the same reaction was performed without SuperScript III.

### ChIP-Nexus

ChIP-Nexus was prepared essentially as described (Shukla et al., 2018). Cells were grown in 4X-YES at 32°C. Briefly, cell pellets corresponding to 7.5×10^8^ cells were lysed by four 1 minute cycles of bead beating in 500 µl of lysis buffer (50 mM HEPES-KOH, pH 7.5, 140 mM NaCl, 1 mM EDTA, 1% Triton X-100, 0.1% sodium deoxycholate). Insoluble chromatin fraction was isolated by centrifugation at 6000×g. The pellet was washed with 1 ml lysis buffer and gently resuspended in 300 µl lysis buffer containing 0.2% SDS and sheared by sonication with Bioruptor (Diagenode) for 30 minutes (30s On, 30s off at high setting). 900 µl of lysis buffer (without SDS) was added and samples were clarified by centrifugation at 17000×g for 20 minutes. Supernatants were used for ChIP. Supernatants were incubated with 6 µl anti-HA (12CA5, in-house preparation) or 6 µl anti-T7 (69522, Millipore) and protein G-dynabeads (ThermoFisher Scientific) for ChIP.

ChIP-Nexus libraries were prepared essentially as described (He et al., 2015). DNA was end repaired using T4 DNA polymerase (NEB, M0203S), DNA polymerase I large fragment (NEB, M0210S) and T4 polynucleotide kinase (NEB, M0201S). A single 3’-A overhang was added using Klenow exo-polymerase. Adapters were ligated and blunted again by Klenow exo-polymerase to fill in the 5’ overhang first and then by T4 DNA polymerase to trim possible 3’ overhangs. Blunted DNA was then sequentially digested by lambda exonuclease (NEB, M0262S) and RecJf (NEB, M0264L). Digested single strand DNA was then eluted, reverse cross-linked and phenol-chloroform extracted. Fragments were then self-circularized by Circligase (Epicentre, CL4111K). An oligonucleotide was hybridized to circularized single DNA for subsequent BamHI digestion in order to linearize the DNA. This linearized single strand DNA was then PCR-amplified using adapter sequences and libraries were purified and size selected using Ampure XP beads. The libraries were sequenced following Illumina HiSeq2500 work flow.

More than 20 million 50bp long single end reads were generated for each sample in this experiment. All reads first 15bp were trimmed, then aligned on S.pombe v2.20 genome build with bowtie2 (Langmead and Salzberg, 2012). Alignment analysis were used by samtools (Li et al., 2009) and bedtools (Quinlan and Hall, 2010). Genome wide enrichment visualization was use deeptools (Ramírez et al., 2014) and IGV browser. ChIP peak were called by MACE 1.2 (Wang et al., 2014).

## DATA AND SOFTWARE AVAILABILITY

The accession number for the sequencing data reported in this paper is GEO: GSE136305. The mass spectrometry proteomics data have been deposited to the ProteomeXchange Consortium via the PRIDE (Vizcaino et al., 2013) partner repository with the dataset identifier PXD015222.

## ACKNOWLEDGEMENTS

We thank members of the Allshire lab for advice and useful suggestions. We thank Dominik Hoelper for critical reading of the manuscript. This research was supported by award of a Darwin Trust of Edinburgh Studentship to P.P.S, a Wellcome Senior Research Fellowship 103139) and Wellcome Instrument grant to J.R. (108504), a Wellcome Principal Research Fellow to R.C.A. (095021; 200885), and core funding for the Wellcome Centre for Cell Biology (203149).

## AUTHOR CONTRIBUTIONS

P.P.S. and R.C.A. jointly conceived the study. P.P.S., M.S., and S.A.W. performed experiments. P.P.S. and P.T. analysed data. T.A. provided guidance in sample preparation for mass spectrometry. C.S. and J.R. performed mass spectrometry and MaxQuant analysis. P.P.S., A.L.P., and R.C.A. wrote the manuscript with input from other authors.

## Supplemental Information

**Table S1.**
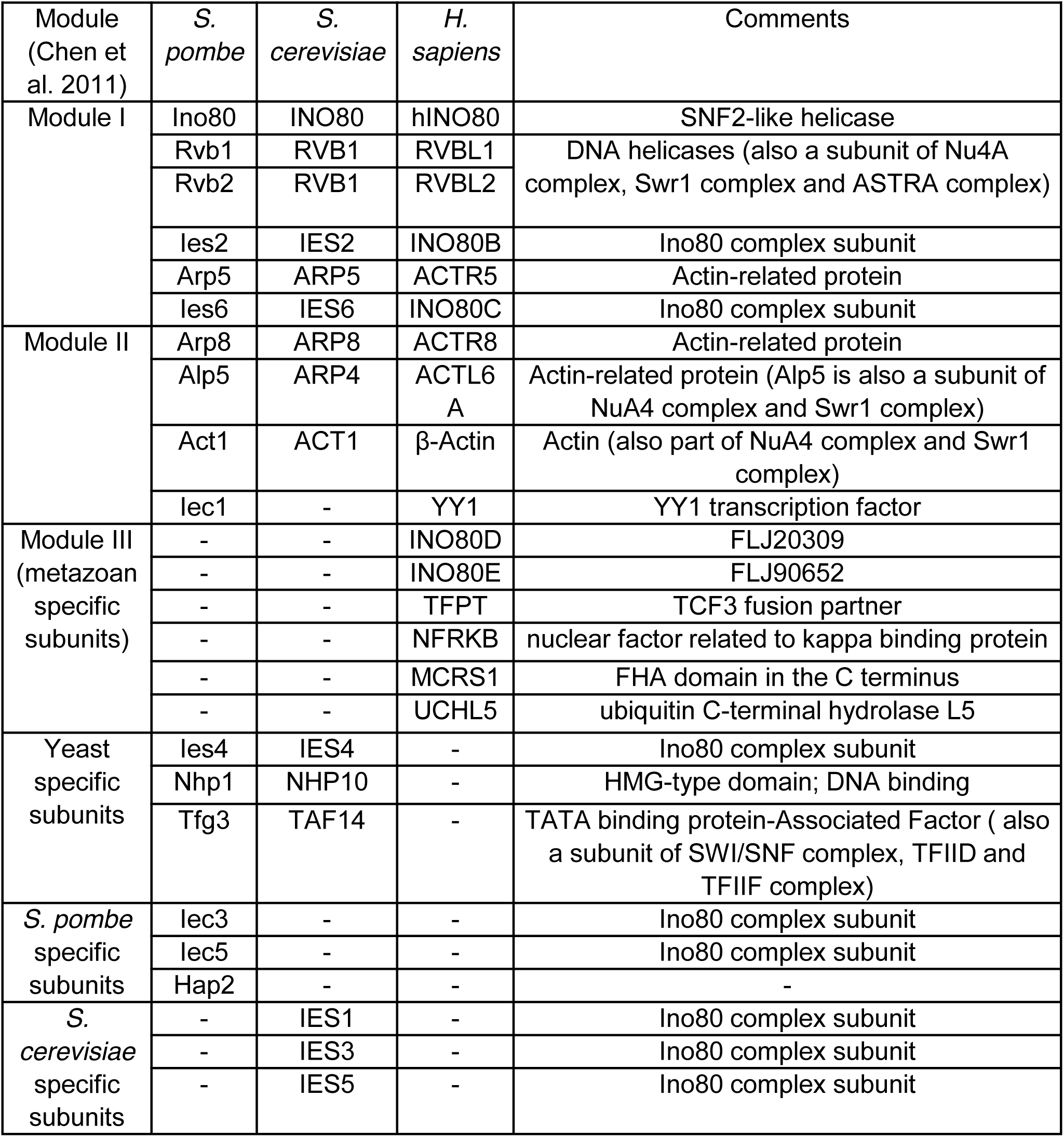
Compositions of *S. pombe, S. cerevisiae* and *H. sapiens* Ino80-related chromatin remodeling complex.

**Table S2.**
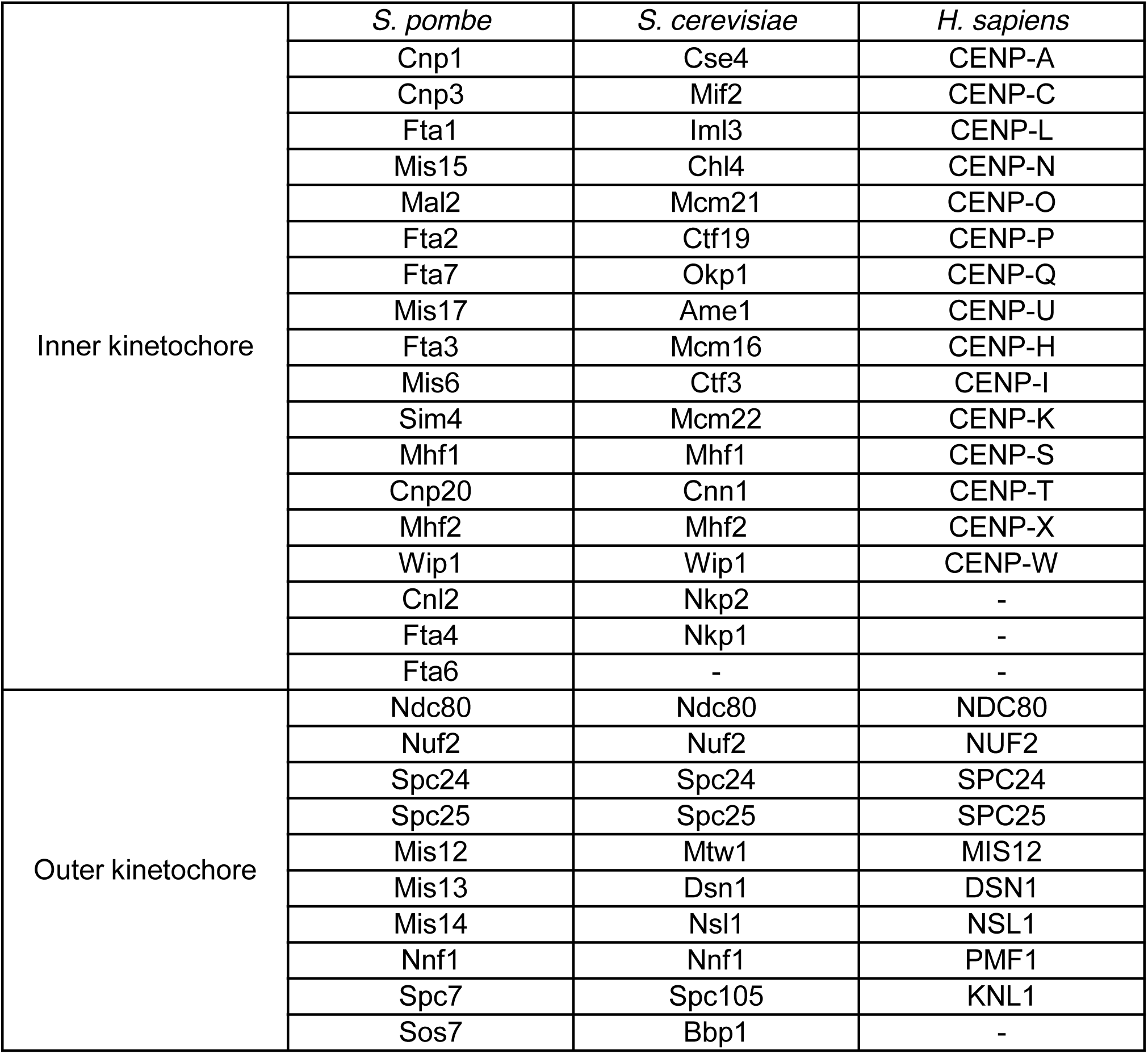
Orthologous *S. pombe, S. cerevisiae* and *H. sapiens* kinetochore proteins.

|  | *S. pombe* | *S. cerevisiae* | *H. sapiens* |
| --- | --- | --- | --- |
| Inner kinetochore | Cnp1 | Cse4 | CENP-A |
|  | Cnp3 | Mif2 | CENP-C |
|  | Fta1 | Iml3 | CENP-L |
|  | Mis15 | Chl4 | CENP-N |
|  | Mal2 | Mcm21 | CENP-O |
|  | Fta2 | Ctf19 | CENP-P |
|  | Fta7 | Okp1 | CENP-Q |
|  | Mis17 | Ame1 | CENP-U |
|  | Fta3 | Mcm16 | CENP-H |
|  | Mis6 | Ctf3 | CENP-I |
|  | Sim4 | Mcm22 | CENP-K |
|  | Mhf1 | Mhf1 | CENP-S |
|  | Cnp20 | Cnn1 | CENP-T |
|  | Mhf2 | Mhf2 | CENP-X |
|  | Wip1 | Wip1 | CENP-W |
|  | Cnl2 | Nkp2 | - |
|  | Fta4 | Nkp1 | - |
|  | Fta6 | - | - |
| Outer kinetochore | Ndc80 | Ndc80 | NDC80 |
|  | Nuf2 | Nuf2 | NUF2 |
|  | Spc24 | Spc24 | SPC24 |
|  | Spc25 | Spc25 | SPC25 |
|  | Mis12 | Mtw1 | MIS12 |
|  | Mis13 | Dsn1 | DSN1 |
|  | Mis14 | Nsl1 | NSL1 |
|  | Nnf1 | Nnf1 | PMF1 |
|  | Spc7 | Spc105 | KNL1 |
|  | Sos7 | Bbp1 | - |

**Figure S1.**
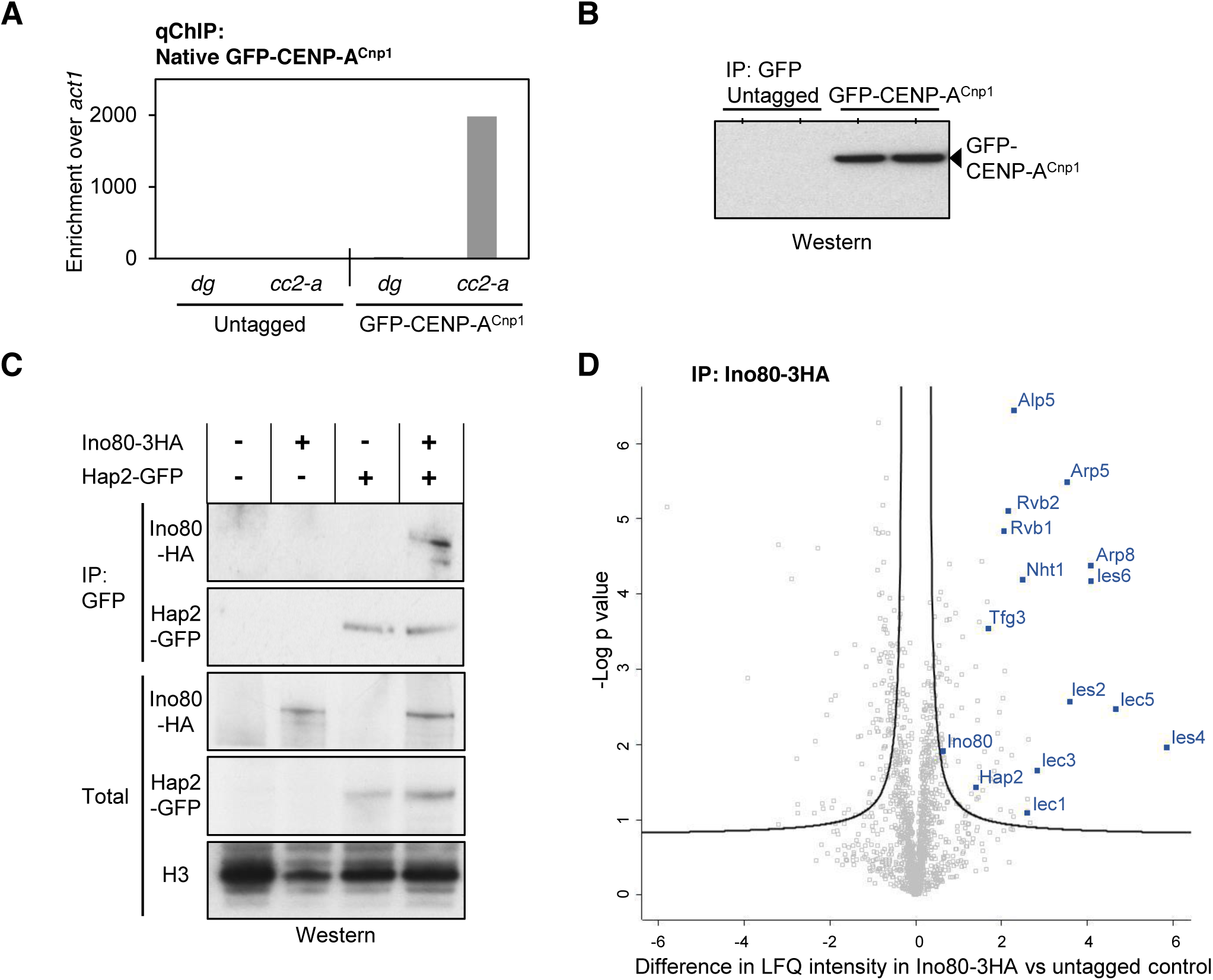
Purification of CENP-A^Cnp1^ and Ino80 complex. (A) Native qChIP for GFP-CENP-A^Cnp1^ at outer repeat heterochromatin (*dg*) and within endogenous centromeres (*cc2-a*). (B) Anti-GFP IP from lysates of cells expressing GFP-CENP-A^Cnp1^ or untagged control cells, followed by anti-GFP western analysis. (C) Western analysis of anti-GFP IPs to enrich Hap2-GFP from lysates of cells expressing Ino80-3HA, Hap2-GFP or both, or untagged tagged control. Western analysis to detect Ino80-3HA or Hap2-GFP in anti-GFP IPs and total extracts. Loading control anti-histone H3. (D) Volcano plot comparing LFQ intensity of proteins enriched in affinity selected Ino80-3HA (anti-HA) versus untagged control. Ino80 complex subunits (Blue).

**Figure S2.**
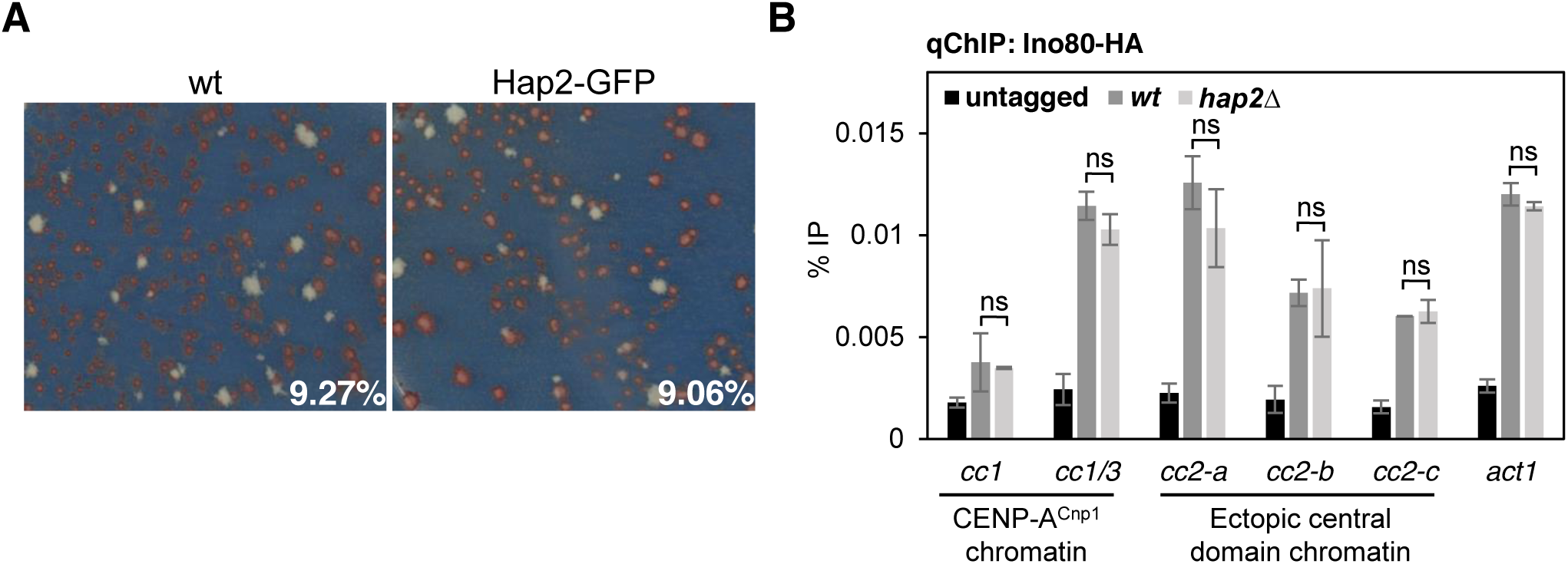
C-terminal tagging of Hap2 is functional Ino80 localization at central domain chromatin is independent of Hap2. (A) *S. pombe* transformed with pHcc2 minichromosome DNA were replica-plated to low adenine non-selective plates. Colonies where CENP-A^Cnp1^ chromatin and functional centromeres have established retain pHcc2 and exhibit a white/pale-pink colour. Those that do not establish centromeres lose pHcc2 and form red colonies. Representative plates of such establishment assay are shown for wild type (n = 539) and Hap2-GFP (n = 342) cells. The percentage of transformants exhibiting functional centromere establishment are indicated. (B) qChIP for Ino80-HA at two locations within endogenous centromeres (*cc1* and cc1/3), three locations within ectopically inserted central domain 2 DNA (*cc2-a*, *cc2-b* and *cc2-c*) and non-centromere locus (*act1^+^*), in wild type and *hap2*Δ. Error bars indicate mean ± SD (n = 3). Significance of the differences observed between wild type and *hap2*Δ was evaluated using Student’s t-test; * *p* < 0.05, ** *p* < 0.005; *** *p* < 0.005 and n.s., not significant.

**Figure S3.**
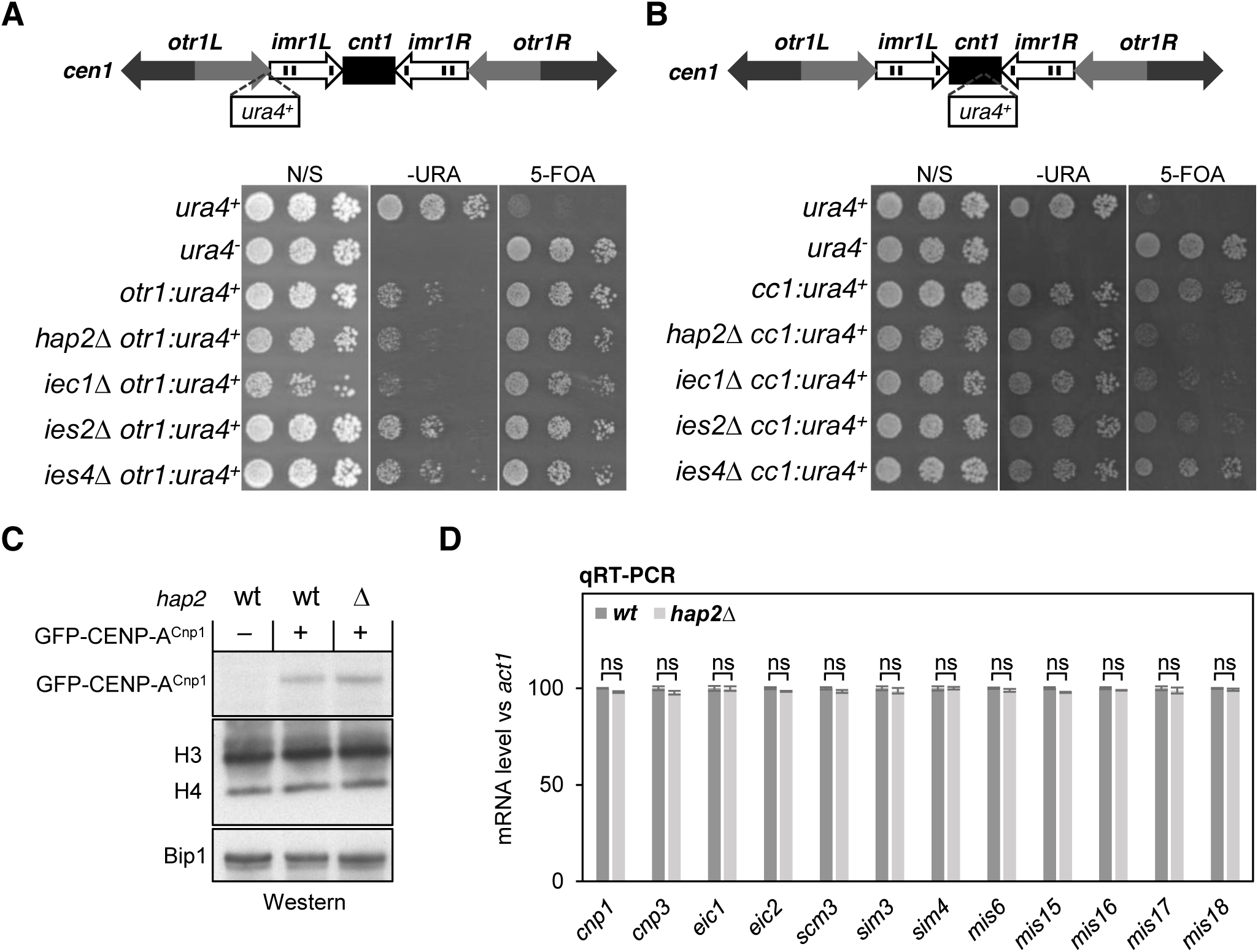
Loss of Hap2 does not affect cellular levels of CENP-A^Cnp1^ or mRNA levels of CENP-A^Cnp1^ chromatin-related genes. (A and B) Growth assay for at *cnt1-otr1:ura4^+^* (A) or *cnt1-cc1:ura4^+^* (B) in cells lacking the indicated Ino80 subunits. (C) Western analysis of GFP-CENP-A^Cnp1^, histone H3, histone H4 and Bip1 (loading control) in whole cell lysates of wild-type and *hap2*Δ cells expressing GFP-CENP-A^Cnp1^ or untagged CENP-A^Cnp1^. (D) qRT-PCR analysis to measure expression levels of the indicated genes which are known to affect CENP-A^Cnp1^ loading at fission yeast centromeres when mutated. Transcript levels are shown relative to the *act1^+^* gene (n = 3). Significance of the differences observed between wild type and *hap2*Δ was evaluated using Student’s t-test; * *p* < 0.05, ** *p* < 0.005; *** *p* < 0.005 and n.s., not significant.

**Figure S4.**
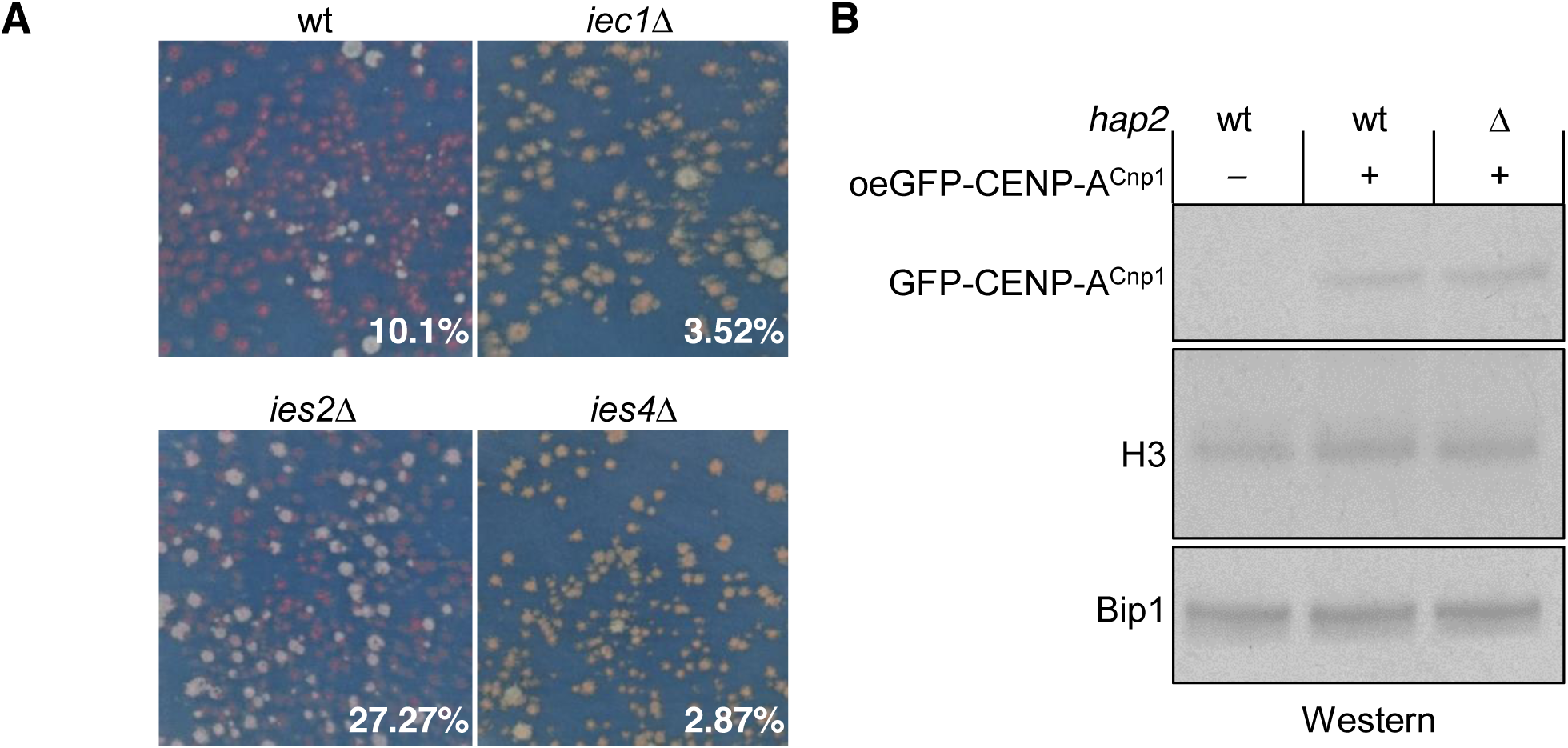
Cells lacking different Ino80 subunits show variable centromere establishment frequency Overexpression of CENP-A^Cnp1^ is unaffected by the loss of Hap2. (A) *S. pombe* transformants containing pHcc2 minichromosome plasmids were replica-plated to low adenine non-selective plates: colonies retaining pHcc2 minichromosome plasmid have established centromeres are white/pale-pink, those that lose it are red. Representative plate showing established and non-established colonies and establishment frequency in wild type (n = 515), *iec1*Δ (n = 795), *ies2*Δ (n = 660) and *ies4*Δ (n = 802). (B) Western analysis of GFP-CENP-A^Cnp1^, histone H3, histone H4 and Bip1 (loading control) in whole cell lysates of wild-type and *hap2*Δ cells over-expressing GFP-CENP-A^Cnp1^ or untagged CENP-A^Cnp1^.

**Figure S5.**
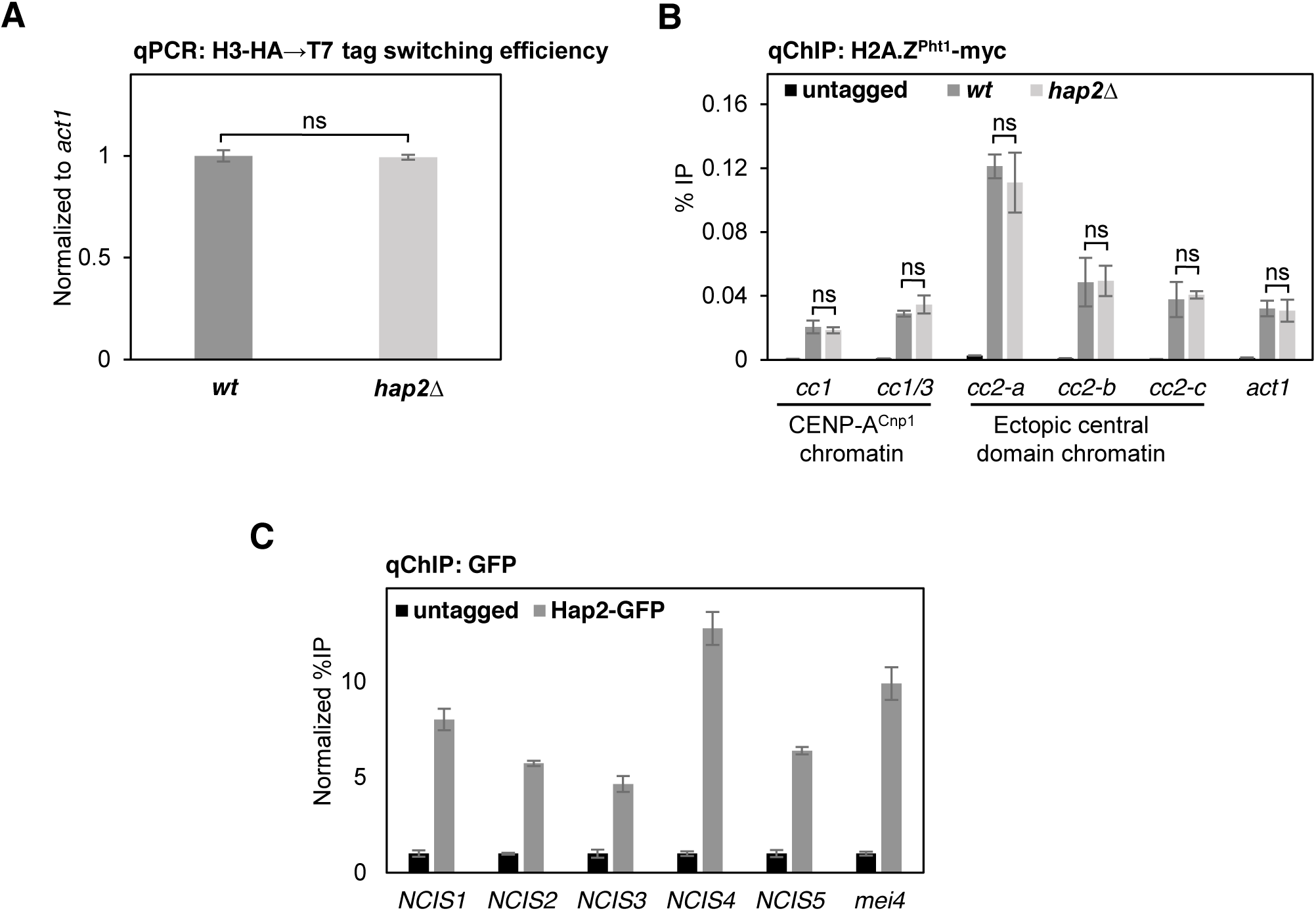
H3-HA→T7 switching efficiency is not affected in cells lacking Hap2 H2A.Z^Pht1^ localization at central domain chromatin is independent of Hap2 Hap2-GFP associates with non-centromeric CENP-A^Cnp1^ islands. (A) qPCR for H3-HA→T7 switching efficiency of H3 tagged with HA and switched to T7 tag by β-estradiol induced Cre/loxP mediated recombination during *cdc25-22*/G2 block in 2 hours. Error bars indicate mean ± SD (n = 3). (B) qChIP for H2A.Z^Pht1^ at two locations within endogenous centromeres (*cc1* and cc1/3), three locations within ectopically inserted central domain 2 DNA (*cc2-a*, *cc2-b* and *cc2-c*) and non-centromere locus (*act1^+^*), in wild type and *hap2*Δ. Error bars indicate mean ± SD (n = 3). (C) qChIP for Hap2-GFP at ectopic CENP-A^Cnp1^ islands (*NCIS1*, *2*, *3* and *4*) and meiotic-specific gene (*mei4^+^*) in untagged and Hap2-GFP. Error bars indicate mean ± SD (n = 3). Error bars indicate mean ± SD (n = 3). Significance of the differences observed between wild type and *hap2*Δ was evaluated using Student’s t-test in A; * *p* < 0.05, ** *p* < 0.005; *** *p* < 0.005 and n.s., not significant.

**Figure S6.**
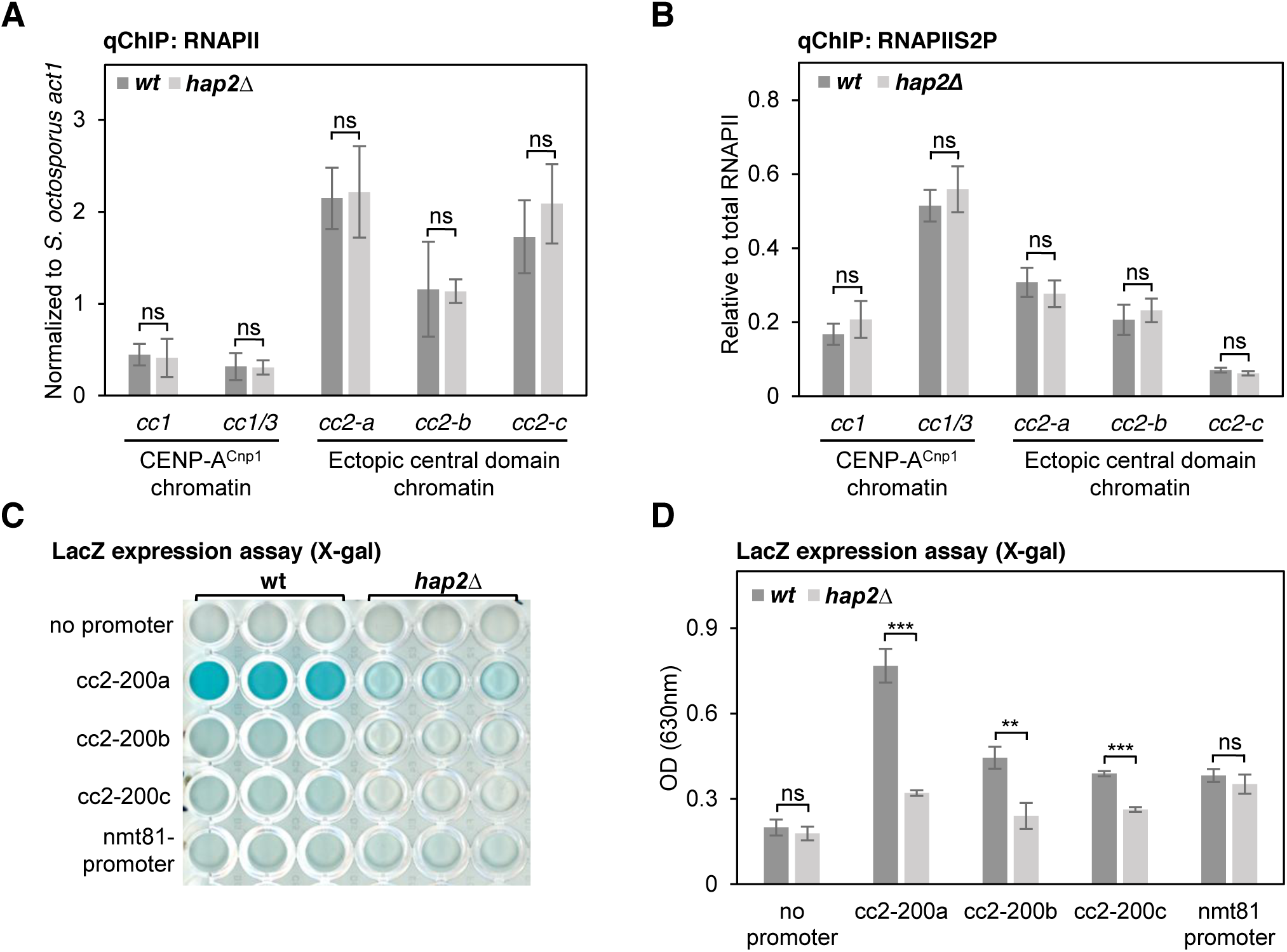
Transcription from centromeric DNA is reduced in the absence of Hap2. (A and B) qChIP for total RNAPII and RNAPIIS2P at two locations within endogenous centromeres (*cc1* and cc1/3) and three locations within ectopically-inserted central core 2 DNA (*cc2-a*, *cc2-b* and *cc2-c*) in wild type and *hap2*Δ in G3 (T140). Error bars indicate mean ± SD (n = 3). (C) Analysis of promoter activity from cc2 fragments (cc2-200a, cc2-200b and cc2-200c). The level of LacZ expression was assessed by measuring absorbance at 630 nm of cell lysates incubated with X-gal. nmt81: positive control with nmt81 promoter. (D) Quantification of levels of LacZ expression as assessed by measuring absorbance at 630 nm in samples shown in C (n = 3). Error bars indicate mean ± SD (n = 3). Significance of the differences observed between wild type and *hap2*Δ was evaluated using Student’s t-test in A, B and D; * *p* < 0.05, ** *p* < 0.005; *** *p* < 0.005 and n.s., not significant.

**Table S3.** List of strains used in the study.

| Strain no. | Genotype | Reference |
| --- | --- | --- |
| 1646 | *h- ade6-210 arg3-D4 his3-D1 leu1-32 ura4-D18* | Lab stock |
| 4841 | *h- otr1R Sph::ura4 leu1-32 ura4DSE his3D1 arg3D4 ade6-210* | Allshire et al., 1995 |
| A2549 | *h- leu1-32 ura4-D18 lys1 GFP-cnp1-NAT* | Takayama et al., 2008 |
| A7373 | *h- ade6-704-HYGMX6 his3-D1 leu1-32 ura4-DSE/D18? arg3? cc2D6kb:cc1* | Catania et al., 2015 |
| A7377 | *clr4::NAT ade6-704-HYGMX6 his3-D1 leu1-32 ura4-DSE/D18? arg3? cc2D6kb:cc1* | Catania et al., 2015 |
| A7408 | *h- cc2D6kb:cc1 ars1:nmt41-GFP-cnp1-NAT ade6-704-HYGMX6 his3-D1 leu1-32 ura4-DSE/D18* | Catania et al., 2015 |
| A8240 | *h- kanMX6-Pcnp1-mEGFP-cnp1 ade6-M210 leu1-32 ura4-D18* | JW2595. Gift from Jian-qiu Wu. |
| A8545 | *h+ ade6-210 arg3-D4 his3-D1 leu1-32 ura4-D18 cnp1:cnp1-loxP-HA-HYG-loxP-T7* | This study |
| A8548 | *h+ ade6-210 arg3-D4 his3-D1 leu1-32 ura4-D18 cnp1:cnp1-loxP-T7-HYG-loxP-HA* | This study |
| A8985 | *Ura4-DSE-sup3e-cc2-ura4+ cc2D6kb:cc1 ade6-704-HYG ura4-DSE/D18?* | Lab stock |
| B0324 | *cc2Δ6Kb:cc1 Ura4-DSE-sup3e-cc2-ura4+ H3.2-lox-HA-HYG-loxT7 ars1:pRAD11-Cre-EBD-LEU2+ cdc25-22 ade6-704-NAT ura4D18/DSE leu1-32* | Shukla et al., 2018 |
| B1700 | *iec1::Kan TM1::ura ade6-210/216 ura-D18 leu1-35* | This study |
| B1713 | *ies2::Kan TM1::ura ade6-210/216 ura-D18 leu1-48* | This study |
| B1715 | *ies4::Kan TM1::ura ade6-210/216 ura-D18 leu1-50* | This study |
| B1718 | *hap2::Kan TM1::ura ade6-210/216 ura-D18 leu1-53* | This study |
| B1819 | *h+ ade6-M210/M216 ura4-D18 leu1-32 hap2::KanMX6* | This study |
| B1863 | *cc2D6kb:cc1 ars1:nmt41-GFP-cnp1-NAT ade6-704-HYGMX6 leu1-32 ura4-DSE/D18 iec1::KanMX6* | This study |
| B1884 | *cc2D6kb:cc1 ars1:nmt41-GFP-cnp1-NAT ade6-704-HYGMX6 leu1-32 ura4-DSE/D18 ies2::KanMX6* | This study |
| B1887 | *cc2D6kb:cc1 ars1:nmt41-GFP-cnp1-NAT ade6-704-HYGMX6 leu1-32 ura4-DSE/D18 ies4::KanMX6* | This study |
| B1890 | *cc2D6kb:cc1 ars1:nmt41-GFP-cnp1-NAT ade6-704-HYGMX6 leu1-32 ura4-DSE/D18 hap2::KanMX6* | This study |
| B1929 | *ade6-704-HYGMX6 leu1-32 cc2D6kb:cc1 hap2Δ::KanMX6* | This study |
| B2132 | *h+ ura4-D18 leu1-32 GFP-cnp1-NAT hap2Δ::Kan* | This study |
| B2871 | *hap2::Kan cc2?6Kb:cc1 Ura4-DSE-sup3e-cc2-ura4+ H3.2-lox-HA-HYG-loxT7 ars1:pRAD11-Cre-EDB-LEU2+ cdc25-22 ade6-704-NAT ura4D18/DSE leu1-32* | This study |
| B3109 | *h- hap2-GFP-Kan ade6-210 arg3-D4 his3-D1 leu1-32 ura4-D18* | This study |
| B3646 | *leu1-32 ura4-D18 Ino80-HA-NAT* | This study |

**Table S3. List of strains used in the study (continued)**
| Strain no. | Genotype | Reference |
| --- | --- | --- |
| B3993 | *h- pht1-myc-KanMX ura4-D18/DS-E*<br>*Ura4-DSE-sup3e-cc2-ura4+ cc2D6kb:cc1 ade6-704-HYG* | This study |
| B4079 | *Ino80-3HA-Nat ura4-D18/DS-E?*<br>*Ura4-DSE-sup3e-cc2-ura4+ cc2D6kb:cc1 ade6-704-HYG* | This study |
| B3854 | *iec1Δ::Kan otr1R Sph::ura4 ura4-D18/DSE leu1-32*<br>*ade6-M210/M216* | This study |
| B3856 | *ies2Δ::Kan otr1R Sph::ura4 ura4-D18/DSE leu1-32*<br>*ade6-M210/M216* | This study |
| B3858 | *ies4Δ::Kan otr1R Sph::ura4 ura4-D18/DSE leu1-32*<br>*ade6-M210/M216* | This study |
| B3860 | *hap2Δ::Kan otr1R Sph::ura4 ura4-D18/DSE leu1-32*<br>*ade6-M210/M216* | This study |
| B4282 | *hap2-GFP-KanMX ura4-D18/DS-E cc2D6kb:cc1*<br>*ade6-704-HYG* | This study |
| B4283 | *hap2-GFP-KanMX ura4-D18/DS-E*<br>*Ura4-DSE-sup3e-cc2-ura4+ cc2D6kb:cc1 ade6-704-HYG* | This study |
| B4285 | *hap2-GFP-KanMX ura4-D18/DS-E cdc25-22*<br>*Ura4-DSE-sup3e-cc2-ura4+ cc2D6kb:cc1 ade6-704-HYG* | This study |
| B4286 | *hap2Δ::KanMX Ino80-3HA-Nat ura4-D18/DS-E*<br>*Ura4-DSE-sup3e-cc2-ura4+ cc2D6kb:cc1 ade6-704-HYG* | This study |
| B4287 | *hap2Δ::Kan pht1-myc-KanMX ura4-D18/DS-E*<br>*Ura4-DSE-sup3e-cc2-ura4+ cc2D6kb:cc1 ade6-704-HYG* | This study |

**Table S4.** List of plasmids used in the study.

| Plasmid name | Reference |
| --- | --- |
| pHcc2 | Folco et al., 2008 |
| pcc2 | Folco et al., 2008 |
| pLacZ (no promoter) | Catania et al., 2015 |
| pLacZ-POP-N13-200bp (cc2-200a) | Catania et al., 2015 |
| pOP-1c-LacZ-200bp (cc2-200b) | Catania et al., 2015 |
| pLM-14A-LacZ-200bp (cc2-200c) | Catania et al., 2015 |
| pREP81X-LacZ (nmt81 promoter) | Catania et al., 2015 |

**Table S5.**
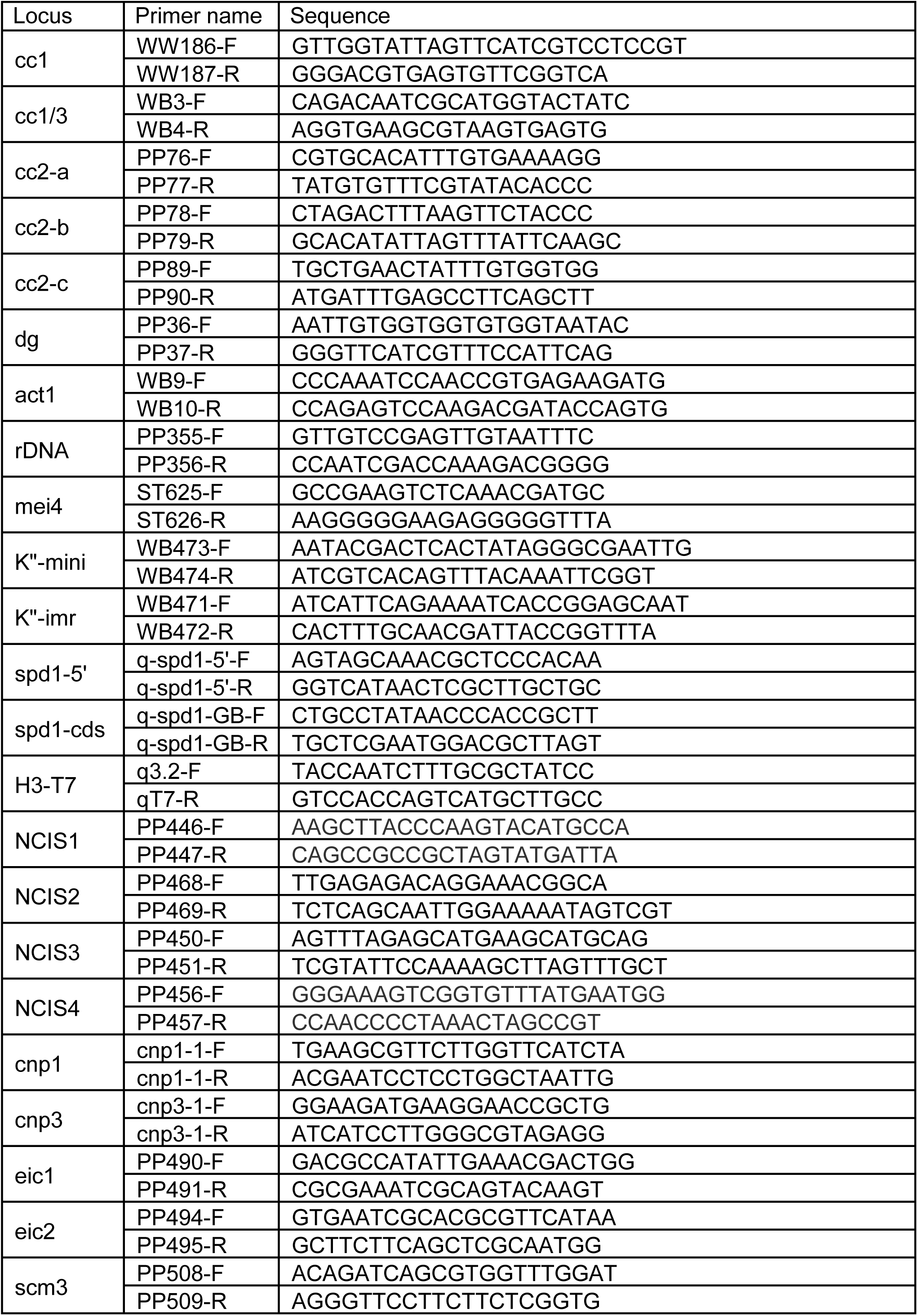
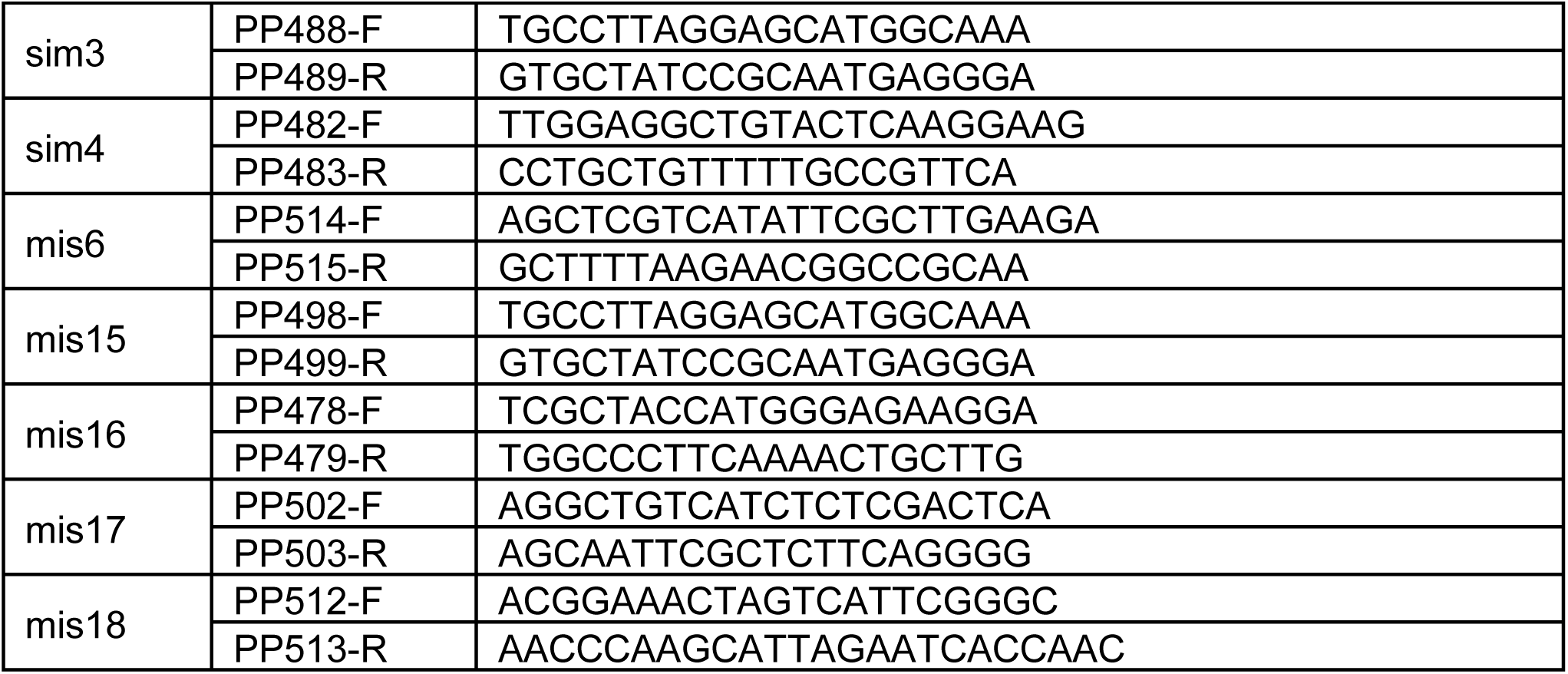
List of primers used in the study.

